# Identifying Neuroimaging Biomarkers of Resilience and Vulnerability to Chronic Stress in An Animal Model: An Exploratory Analysis

**DOI:** 10.1101/2025.08.05.667822

**Authors:** Twain Dai, Shannon Dee Algar, Michael Small, Andrew Zalesky, Jennifer Rodger

## Abstract

Stress is a main contributor to mood disorders, with individuals displaying great heterogeneity in response to stressful life events and adversity. Identifying biomarkers of vulnerability and resilience to stress would facilitate a prevention-based approach to mental illness that benefits individuals and reduces healthcare costs. The present study adopted a multivariate machine-learning approach to track neuroimaging biomarkers predictive of resilience and vulnerability following a chronic restraint stress (CRS) model. 96 male Sprague-Dawley rats underwent two sessions of MRI and behavioral tests, before and after CRS. Resilience and vulnerability to CRS were assessed with elevated plus maze and forced swimming tests. Hierarchical clustering was applied to construct brain networks. Partial correlation was used to compute network connectivity. Repeated nested cross validation with a support vector machine was employed to identify rs-fMRI biomarkers predictive of resilience and vulnerability following CRS. No strong group effect size of behavioral changes following CRS was observed within the same animals, suggesting the presence of resilient and vulnerable subgroups. Although the average model performance was modest (area under the receiver operating characteristic curve: 0.3 ~ 0.67), baseline functional connectivity across cerebellum, brainstem, striatum, prefrontal and salience-orbitofrontal regions, as well as functional alteration across hippocampus, striatum, prefrontal regions, auditory thalamus, cerebellum, inferior colliculi and brainstem were identified as stable features. The present study is the first to identify connectome-based neuroimaging biomarkers predictive of resilience and vulnerability using an animal model. The results may provide insights into neuroimaging biomarkers to aid diagnosis and prevention of mood disorders in humans.

## Introduction

Depression is the second-leading cause of global disability (1), and is characterized by a range of subjective symptoms, including depressed mood, anxiety and psychomotor disturbance. Stress is considered to be one of the main contributing factors to depression (2). But individuals display great heterogeneity in their response to stressful life events and adversity. Some individuals are susceptible to developing depression and/or anxiety in response to stress, while others remain resilient and essentially symptom free. Identifying objective biomarkers of vulnerability and resilience to stress would facilitate a prevention-based approach to depression that benefits individuals and reduces healthcare costs.

Animal models of chronic stress have been widely applied to examine neurobiological mechanisms that shape vulnerability and resilience to stress (3). They facilitate the delivery of controlled interventions that induce behavioral changes considered to model aspects of depression in humans. Similar to humans, emerging evidence reveals a remarkable inter-individual and inter-strain variability in animals’ responses to chronic stress (2-7). Anxiety-like and depression-like behaviors are not exhibited in all animals undergoing chronic stress paradigms, with susceptible and resilient subgroups identified. Resilience or vulnerability to chronic stress in behaviors is found to correlate with distinct regulation of neural and molecular markers. For example, the probability of gamma-aminobutyric acid (GABA) release and frequency of spontaneous inhibitory postsynaptic currents in granule cells of the dentate gyrus are both lower in stress-susceptible than stress-resilient rats (8). Moreover, c-Fos expression in the amygdala, medial habenula, lateral orbital, ventral orbital, and infralimbic cortex is differentially regulated in stress-resilient and stress-susceptible rats (9). In addition, stress-susceptible and stress-resilient rats differ in hypothalamic-pituitary-adrenal (HPA) axis function, with vulnerable male rats displaying upregulation of glucocorticoid receptor in the ventral hippocampus (10). However, these neurobiological markers are obtained post-mortem, precluding the identification of neurobiological features present at baseline or in healthy animals that might predict the separation of vulnerable and resilient subgroups following stress intervention.

Resting-state functional magnetic resonance imaging (rs-fMRI) is a powerful and translationally relevant tool to longitudinally trace biomarkers that predict vulnerability and resilience to stress. It provides a non-invasive way to investigate resting-state functional organization of the brain in both humans and non-human living animals. The resting-state organization in humans is generally referred to as resting-state network (RSN). But RSNs are not unique to humans, homologous functional organization has been observed in the rodent brain (11, 12). For instance, the cingulate and retrosplenial cortex, as core hubs of DMN in humans, have also been identified to form DMN-like functional organization in rodents (12, 13). Moreover, a recent study has shown that the RSN alteration following chronic stress in mice shares similarities with that in human depression, indicating that animal models of chronic stress are translatable to human depression (13). Interestingly, new evidence from rs-fMRI studies reveal that there are high individual differences in the functional connectivity of the rodent brain following chronic stress (12, 14). For example, individual variability in the functional connectivity within DMN-like networks following chronic stress in 96 male rats was observed in our previous study (12). The individual differences may offer a gateway to discovering neuroimaging biomarkers of resilience and vulnerability to chronic stress. This will further provide insights into potential biomarkers for humans that could be used to predict vulnerability to stress and prevent the development of depression because of the shared pathways in depression and stress response across species (15).

However, current research in biomarkers of vulnerability and resilience to stress has relied heavily on univariate group analysis (8-10, 14), which overlooks the individual variability and highly interconnected nature of the brain. It is unlikely that one single region or functional connection can explain all aspects of dysfunctions in the depressed brain. To overcome this limitation, multivariate machine learning approaches, such as feature selection, classification and predictive modelling, have been proposed to inspect the highly dimensional connectome of the whole brain simultaneously and identify connectome patterns specific to individuals (16). Surprisingly, no study so far has applied multivariate machine-learning to model the functional connectivity of the brain and behavior to identify rs-fMRI biomarkers of resilience and vulnerability in rodent models of depression. The present study aimed to identify rs-fMRI biomarkers predictive of vulnerability and resilience in a rodent model of depression, using a multivariate machine-learning approach.

## Methods and Materials

### Ethics Statement

All experimental procedures adhered to the ethics guideline of the University of Western Australia Animal Ethics Committee (RA/3/100/1640) and the National Health and Medical Research Council’s Australian code for the care and use of animals for scientific purposes.

### Study Design and Procedures

The data analyzed in the present study was a combined MRI and behavioral dataset of four rodent cohorts from previous experiments conducted by our lab in 2019 and 2020. All animals were sourced from the Animal Resources Centre (Canning Vale, Western Australia). Animals were housed in pairs under a standard 12-hour light-dark cycle with *ad libitum* food and water, in a temperature-controlled environment located at UWA’s Animal Care Unit, M block building (Nedlands, WA).

96 male Sprague-Dawley rats (aged 6 ~ 7 weeks and weight ranged 150 - 250 g on arrival) underwent a chronic stress model - chronic restraint stress (CRS) for 13 consecutive days (*Figure 1*). CRS, as the most popular animal model of depression, was used to establish anxiety-like and depression-like behaviors (17). The CRS procedure adhered to the protocol specified in Ulloa *et al*. (17). Briefly, each rat was placed in a transparent acrylic tube to restrict their free movement for 2.5 hours daily. Following CRS, rats were returned to their home cages.

**Figure 1.**
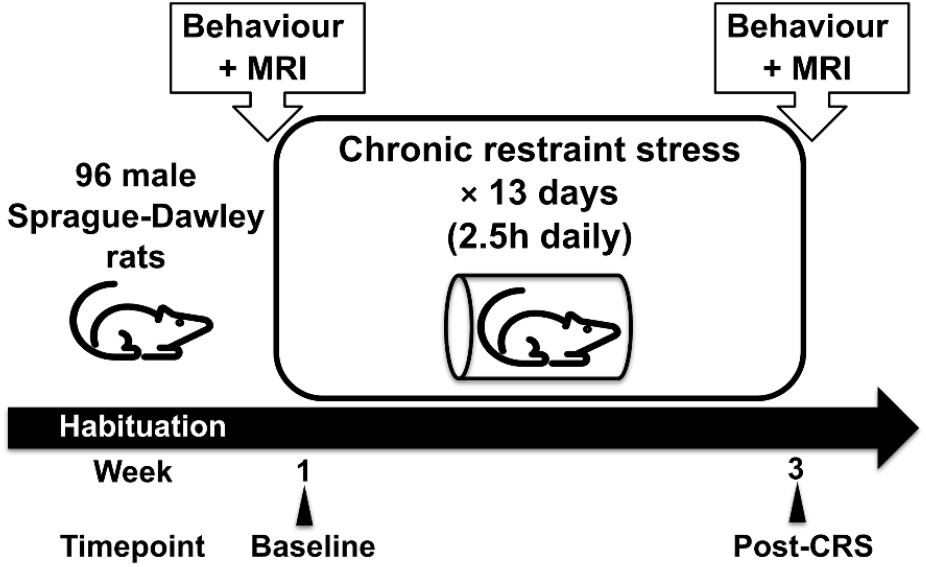
Study design and timeline. All animals were habituated to the new environment for one week after their arrival and prior to experiments. Behavioral tests and MRI scan were conducted before and after 13 consecutive days of CRS.

Behavioral tests included elevated plus maze (EPM) test and forced swimming test (FST). These tests were carried out at baseline (three and four days before the first CRS procedure) and post-CRS (the next two days following the final CRS procedure). EPM and FST tests were conducted following the protocols described in Walf and Frye (18) and Slattery and Cryan (19) to evaluate anxiety-related and stress coping behaviors, respectively. Further description of the protocol can be found in Seewoo *et al*. (20). Each behavioral test session was recorded using a GoPro HERO7 camera.

MRI scans were conducted after the behavioral tests, at both baseline and post-CRS. Animal anesthesia and MRI acquisition protocols were described and discussed thoroughly in the previous publications (12, 20). Brief description can also be found in *Supplementary Text*.

Behavioral tests and MRI scans were conducted in a consistent order of EPM, FST and MRI on different days, for all animals.

### Data Processing

Most of the EPM video and MRI processing were remotely operated on a heavy computing desktop with two Nvidia-T4 GPUs and 225 GB RAM (21), unless otherwise specified.

#### FST Manual Annotation

A total of 158 FST recordings from 79 animals (46 from Hennessy *et al*. (22) and Seewoo *et al*. (20), and 33 unpublished) were acquired at baseline and post-CRS. The recordings from the other 17 rats were either corrupted or could not be saved due to technical failure. FST videos were blinded and manually annotated by researchers. the Floating and active motor behaviors were scored using a time-sampling approach (19). Active motor behaviors included swimming and climbing. More descriptions of the manual annotation can be found in *Supplementary Text*. Activity score (23) was used to measure animals’ overall activity, calculated by subtracting the total counts of floating behavior from active motor behaviors. A negative activity score indicated a passive coping behavior (more time spent in floating), while a positive score suggested an active coping behavior (more time spent in active motor behaviors). Moreover, an activity score of zero indicated neutral coping behavior (equal time spent in floating and active behaviors). Non-parametric estimation statistics based on 5000 bootstrap resampling were performed using dabestr (24) to assess changes in the activity score before and after CRS within the same animals. Paired Cohen’s d (standardized mean difference) was used to measure the effect size of this behavioral change. Results were reported with an estimation plot, Cohen’s d and 95% confidence intervals (CI; 24).

The operational criteria for defining an animal as resilient or vulnerable to developing passive coping behavior following CRS were based on changes in activity scores across the two test sessions. Animals presenting passive coping behavior at baseline (N = 14) were excluded from the classification. Among the remaining animals, those with an increased activity score following CRS were classified as resilient; otherwise, they were defined as vulnerable. An alluvial plot was used to visualize the changes in the coping behavior from baseline to post-CRS within the same animals. A pie chart was employed to depict the proportion and numbers of resilient and vulnerable animals following CRS, evaluated in the FST.

#### EPM Automation

A total of 94 animals completed both EPM sessions, with 188 recordings acquired. One animal fell off the apparatus during EPM test and the recording of the other animal was corrupt. All original EPM video recordings were edited to be five minutes in duration to remove irrelevant (non-test related) footage. The edited videos were then imported into DeepLabCut (version: 2.2.0.6; 25) for further processing. DeepLabCut was used to track 11 body points (Figure 2A) and 12 points of the testing arena (Figure 2B). Main steps involved in the video analysis were as follows: a). Label all the tracking points for 460 randomly selected video frames (20 frames × 23 videos); b). Train a deep neural network/architecture using these 460 labelled frames; c). Apply the trained network to extract the coordinates of tracking points for all edited recordings. For each edited recording, DeepLabCut automatically generated one individual CSV file containing the coordinates of 23 tracking points for all video frames, along with labelling confidence (ranging from 0 - 100%) for each coordinate.

**Figure 2.**
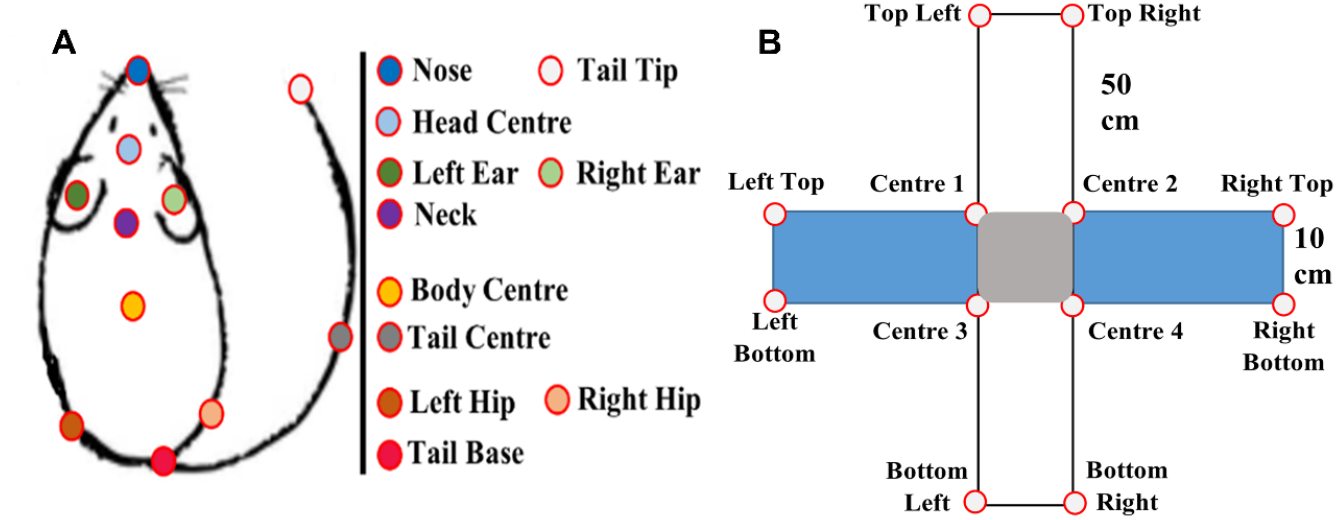
Labels used to train the deep neural network. A). Rat with 11 body points; B). EPM with 12 points. White denotes two oppositely positioned open arms. Blue denotes two closed arms. Grey denotes the center area.

All DeepLabCut coordinate files derived from EPM recordings were imported into RStudio (Windows version: 2023.03.0) and processed with DLCAnalyzer (26) and in-house R scripts (*Supplementary Scripts*). The maze outline for each EPM video was first determined by the median coordinates of each arena point. Rat body points with a likelihood higher than 95% within the maze outline were then used for further processing. The percentage of time spent in closed arms, as a classic measure of anxiety-like behavior, was extracted using the in-house R scripts. Animals spending more than 90%, between 65% and 90%, and less than 65% of the time in closed arms were considered to display high, medium and low anxiety-related behavior, respectively (27). Paired Cohen’s d was used to measure the effect size of changes in the percentage of time spent in closed arms before and after CRS within the same animals. Results were reported with an estimation plot, Cohen’s d and 95% CI (24).

The operational rules for defining an animal as being resilient or vulnerable to developing anxiety-related behaviors following CRS were based on the changes in the percentage of time spent in closed arms across two test sessions. Animals exhibiting high anxiety-like behaviors at baseline (N = 7) were excluded from the classification process. For the remaining animals, those demonstrating consistently low anxiety-like behaviors across both test sessions or showing a reduction in time spent in the closed arms following CRS, were classified as resilient. Conversely, animals not meeting these criteria were defined as vulnerable. An alluvial plot was used to visualize the changes in the percentage of time spent in closed arms from baseline to post-CRS within the same animals. A pie chart was employed to illustrate the proportion and numbers of resilient and vulnerable animals following CRS, evaluated in the EPM.

#### MRI Batch Processing

A total of 192 MRI packages (96 animals × two sessions) were acquired at baseline and post-CRS imaging sessions. The analytic workflow for these packages mostly consisted of common processing steps (12, 28-31). Scripts for executing the workflow can be found in *Supplementary Scripts*, in the order of processing steps. Most of the steps were performed with the Functional MRI of the Brain (FMRIB) Software Library (FSL; version: 6.0.3; 32), unless otherwise specified.

##### A). Pre-Processing

Each data package was first batch processed with an in-house pre-processing pipeline. The pipeline mainly included data organization, bias field correction, brain extraction, denoising, and atlas registration. Detailed pre-processing description can be found in *Supplementary Text*.

##### B). Reproducible Functional Components Identification

Rationale and description for identifying reproducible functional components can be found in the previous publication (12) and *Supplementary Text*. This process identified 217 reproducible functional components/nodes, which were merged to create a group-ICA template of the whole brain gray matter for network modelling.

##### C). Network Modelling to Construct Large-Scale Functional Networks

The group-ICA template was mapped onto de-noised and normalized baseline data (N = 96), to extract timeseries for all nodes for each animal using FSL/fsl_glm (33). These timeseries data were then fed into FSL/FSLNets (version: 0.6) in MATLAB (version: r2019b) to identify spatial clusters for all nodes. A group average cluster hierarchy was constructed using full correlation with Ward’s method. Highly correlated nodes were merged into a large-scale functional network. Natural division and optimal number of networks was determined with Auto-CVI-Tool based on shapes, sizes, densities, and separation distances (SSDS) index (34, 35). The optimal number of networks was determined according to the lowest SSDS index across two to 20 networks.

Within each classified network, brain structures were identified and reported (see *Supplementary Spreadsheet*). Absolute volume and resting-state activity (average Z-score) of each structure were extracted using FSL/fslstats. The percentage volume (absolute volume/anatomical total) of each structure, representing the spatial extent or relative size of resting-state activity, was then calculated. These spatial characteristics were used to classify networks (36). Additionally, the volumes with resting-state activity of brain structures covered by each node within each network were extracted and summarized in *Supplementary Spreadsheet*.

##### D). Functional Connectivity Computation

The classified networks were concatenated using FSL/fslmerge to construct a 4D image. This 4D image was then mapped onto all de-noised and normalized data acquired at baseline and post-CRS imaging session (N = 192), to extract timeseries for all networks for each scan session using FSL/fsl_glm (33). These timeseries data were then fed into FSL/FSLNets (version: 0.6) in MATLAB (version: r2019b) to compute functional connectivity between each pair of networks for each imaging session. Partial correlation was applied to improve mathematical robustness and achieve better estimation of functional connectivity (37-40).

### Brain-Behavior Modelling

Neurobehavioral modelling of CRS resilience and vulnerability evaluated in the FST and EPM were both performed in R 4.4.0. Modelling scripts and corresponding workspace can be found in Supplementary Script and Workspace.RData, respectively. The independent variables for both classification models included baseline network connectivity and the changes in the network connectivity following CRS (post-CRS minus baseline). The modelling process was the same for neurobehavioral modelling with FST and EPM. To build a reasonable model, the present study employed exclusion criterion to exclude animals demonstrating high anxiety-related and passive coping behaviors at baseline, to standardize the sample population. This is similar to human studies excluding individuals with sub-clinical symptoms or high baseline data to avoid bias from this small pool of individuals.

Nested cross-validation was applied to animals classified as resilient and vulnerable following CRS, to construct a classification model using nestedcv (41) and caret (42). To address variations arising from differences in cross-validation splits, the nested cross-validation was repeated 50 times with different random seeds. Each repetition involved nesting two levels of cross-validation loops: an outer loop (five folds) for model evaluation and an inner loop (four folds) for feature and model selection (43). The outer loop partitioned the dataset into five folds, with each fold serving once as the outer testing set and the remaining folds as the outer training set. The outer training set was passed to the inner loop and further split into four inner folds. Within the inner loop, each inner fold was used once as a validation set, with the remaining folds utilized for training. During each step of the inner cross-validation, synthetic minority over-sampling technique (SMOTE) was applied to the training data to address the class imbalance in the animals classified as resilient and vulnerable following CRS (44). Embedded filtering of predictors using Boruta method was also conducted to reduce the dimensionality of the independent variables (45). The hyperparameters and variable set that achieved the best performance on the validation set during the inner cross-validation process, were employed to train a support vector machine (SVM) with a radial basis function kernel on the outer training set. The trained SVM classifier was subsequently assessed using the outer testing set.

Model performance was calculated based on the predictions on the outer testing folds and averaged across 50 repetitions. Summary statistics of model performance metrics, including area under the receiver operating characteristic curve (AUC), balanced accuracy, sensitivity, specificity and F1 score were reported. Variable stability was evaluated using variable importance across models trained and tested within outer loops and total frequency with which each variable was selected in the final models across 50 repetitions. The directionality (either raising the probability of animals being resilient or vulnerable following CRS) of predictors was automatically evaluated using the sign of a t-test when building the SVM classifier with nestedcv (41). Unpaired Cohen’s d was further performed to assess the effect size of predictors between the resilient and vulnerable groups. Results were reported with estimation plots, Cohen’s d and 95% CI (24). The importance, frequency, and directionality of predictors were visualized in divergent plots.

## Results

### FST

The activity score significantly decreased following CRS, at a group level (N = 79; baseline: 24.86 ± 24.76; post-CRS: 18.31 ± 23.78; p = 0.012). However, the effect size of the changes in activity score following CRS was −0.27 (95% CI: −0.58 ~ 0.06, *Figure 3A*), indicating that there was no group effect and high individual variability. 65 out of 79 animals displayed active coping behavior at baseline (*Figure 3B*). Among these 65 animals, changes in overall activity ranged from −56 to 32, with 46 animals classified as vulnerable to developing passive coping behavior following CRS (*Figure 3C*).

**Figure 3.**
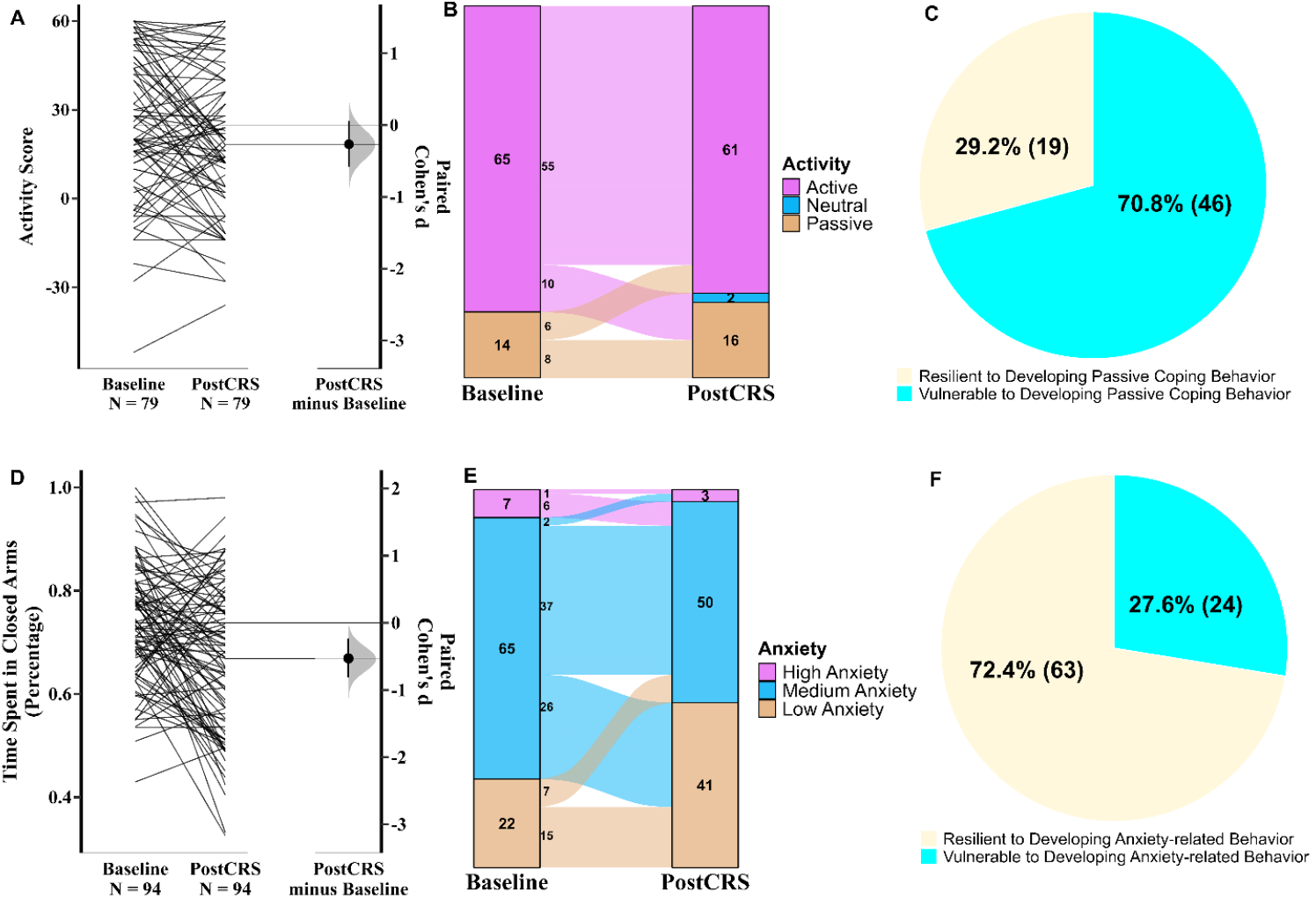
FST and EPM Behaviors. **A). Gardner-Altman estimation plot** showing no group effect size in activity changes in the FST following CRS, with high variability observed. The raw data is plotted on the left axis as a slopegraph, with paired Cohen’s d plotted as a bootstrap sampling distribution on the right axis. Each paired set of observations is connected by a line**; B). Alluvial plot** depicting the changes in the coping behavior from baseline to post-CRS**; C). Pie chart** illustrating the proportion of resilient and vulnerable animals to developing passive coping behavior following CRS; **D). Gardner-Altman estimation plot** showing a medium group effect size in changes of time spent in closed arms in the EPM following CRS, with high variability observed; **E). Alluvial plot** depicting the changes in the anxiety-related behavior from baseline to post-CRS**; F). Pie chart** illustrating the proportion of resilient and vulnerable animals to developing anxiety-related behavior following CRS.

### EPM

The percentage of time spent in closed arms significantly decreased following CRS, at a group level (N = 94; baseline: 0.74 ± 0.11; post-CRS: 0.67 ± 0.11; p = 0.000). But the effect size of the changes in percentage of time spent in closed arms following CRS was −0.529 (95% CI: −0.808 ~ −0.235, *Figure 3D*), indicating a medium group effect and high individual variability. Seven animals displaying high anxiety-like behavior at baseline (*Figure 3E*) were excluded from behavioral classification. Among the remaining 87 animals, changes in percentage of time spent in closed arms spanned from −0.49 to 0.26, with only 24 animals classified as vulnerable to developing anxiety-like behavior following CRS (*Figure 3F*).

### Network Modelling

The data-driven approach - hierarchical clustering merged 217 nodes into 13 large-scale functional networks in the rat brain (*Figure 4, Supplementary Figure S1 & S2*). These networks included salience-orbitofrontal, lateral-temporal cortical, hippocampal-cortical, default mode-somatomotor, spatial attention, corpus striatum-cortical, ventral striatum-cortical, hypothalamus-thalamus, auditory thalamus, (para)flocculus-brainstem and three cerebellum-auditory processing networks. Spatial symmetry was presented in some homologous brain regions within the majority of these networks (detailed description and discussion can be found in the *Supplementary Text)*.

**Figure 4.**
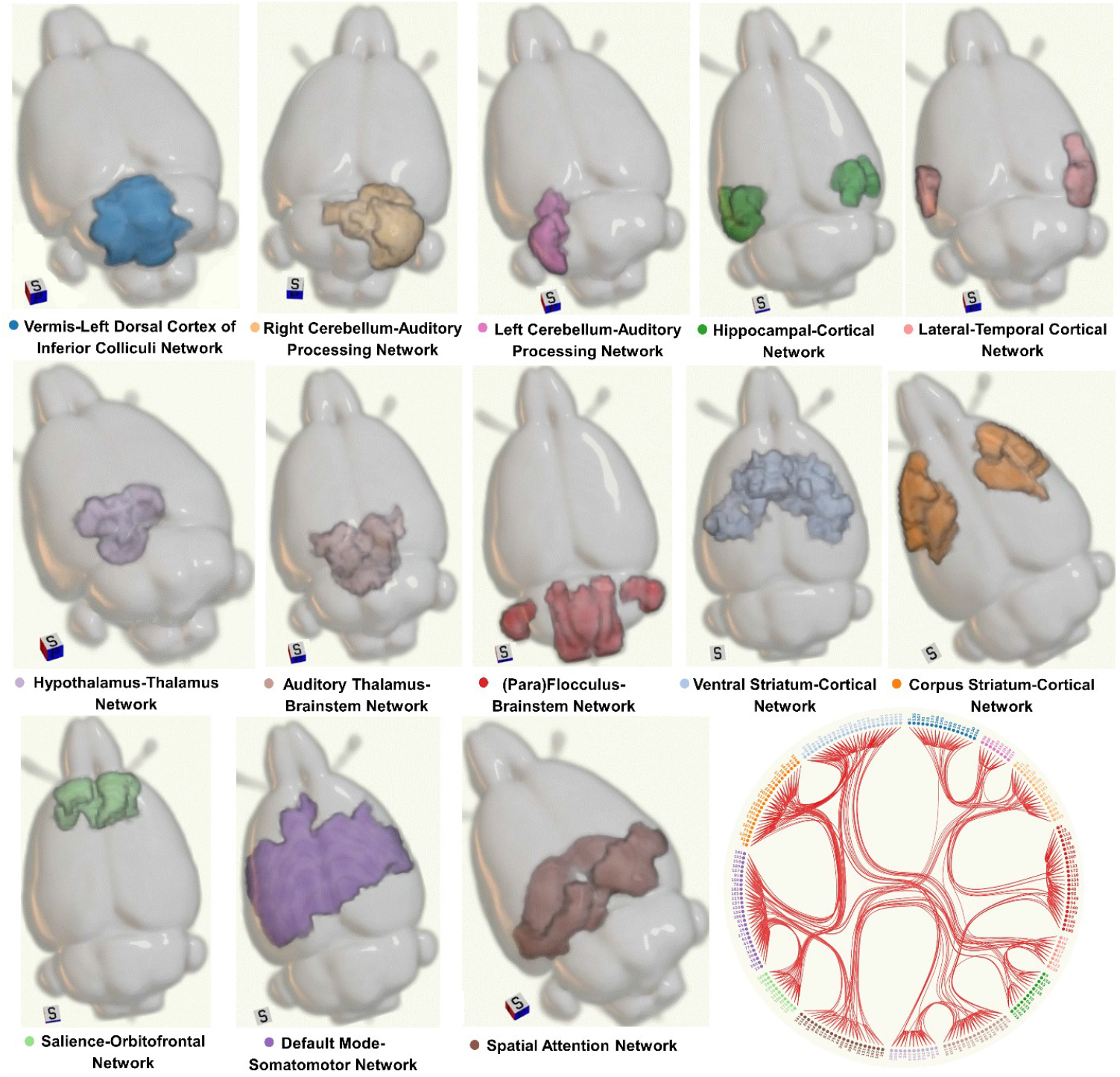
13 Large-Scale Functional Networks. On the bottom right, a chord diagram shows the formation of 13 networks based on hierarchical clustering.

### Neurobehavioral Modelling of CRS Resilience and Vulnerability in FST

Repeated nested cross-validation was conducted on 65 animals when modelling CRS resilience and vulnerability with FST. The model initially incorporated 156 independent variables, including baseline network connectivity 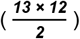, and alteration in the network connectivity 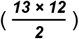. On average, the model demonstrated poor discrimination between resilient animals from vulnerable animals with an average AUC of 0.53 and maximum AUC of 0.67 across 50 repetitions of nested cross-validation (*Table 1*). The F1 score achieved an average score of 0.36, ranging from 0.18 to 0.48. The balanced accuracy spanned from 0.43 to 0.62, with an average of 0.54. The model demonstrated an average sensitivity of 0.39 (range: 0.16 ~ 0.63) and an average specificity of 0.68 (range: 0.54 ~ 0.80).

**Table 1.**
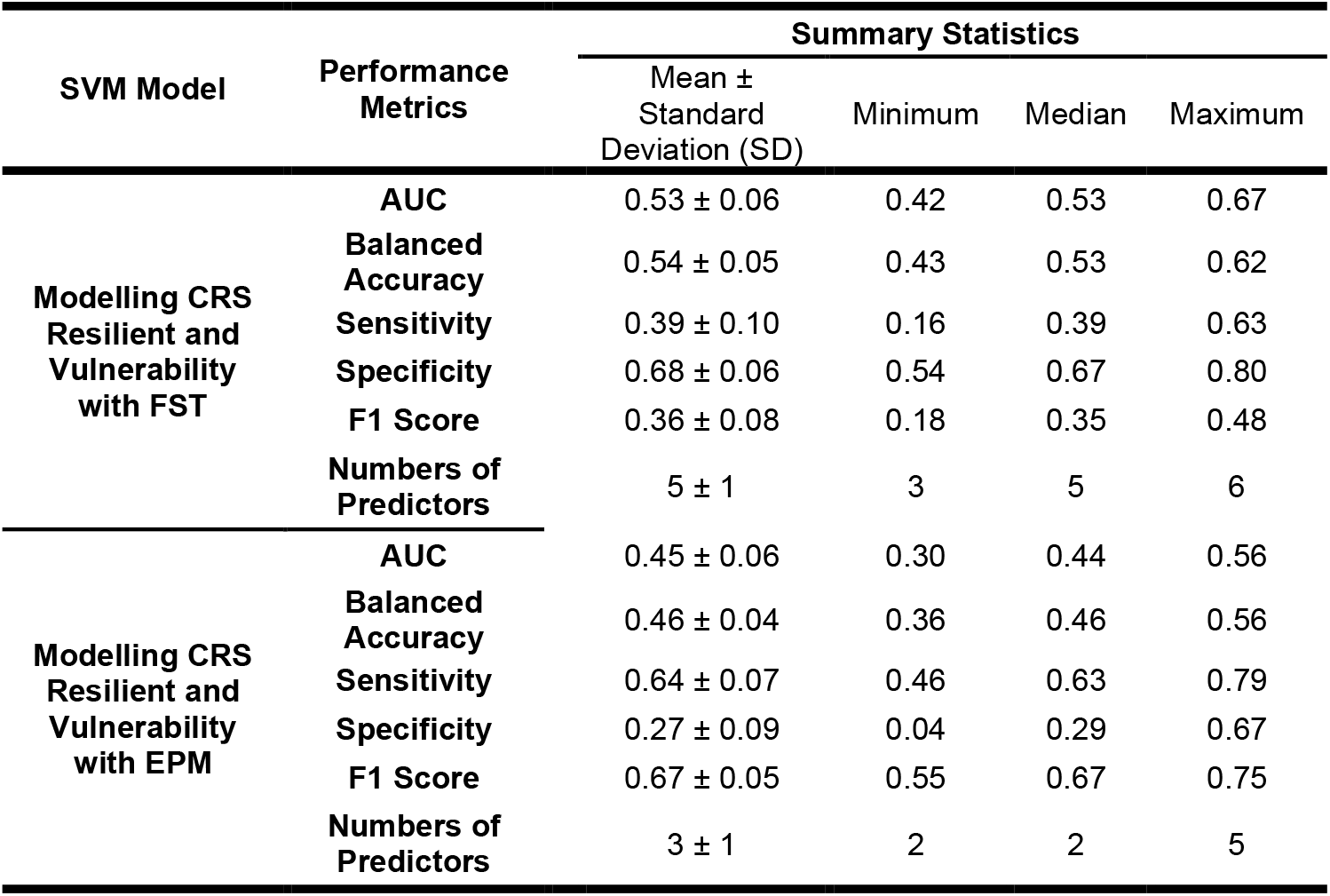
Classification Performance across 50 Repetitions of Nested Cross-Validation.

| SVM Model | Performance Metrics | Summary Statistics |  |  |  |
| --- | --- | --- | --- | --- | --- |
| | | Mean ± Standard Deviation (SD) | Minimum | Median | Maximum |
| Modelling CRS Resilient and Vulnerability with FST | AUC | 0.53 ± 0.06 | 0.42 | 0.53 | 0.67 |
| | Balanced Accuracy | 0.54 ± 0.05 | 0.43 | 0.53 | 0.62 |
| | Sensitivity | 0.39 ± 0.10 | 0.16 | 0.39 | 0.63 |
| | Specificity | 0.68 ± 0.06 | 0.54 | 0.67 | 0.80 |
| | F1 Score | 0.36 ± 0.08 | 0.18 | 0.35 | 0.48 |
| | Numbers of Predictors | 5 ± 1 | 3 | 5 | 6 |
| Modelling CRS Resilient and Vulnerability with EPM | AUC | 0.45 ± 0.06 | 0.30 | 0.44 | 0.56 |
| | Balanced Accuracy | 0.46 ± 0.04 | 0.36 | 0.46 | 0.56 |
| | Sensitivity | 0.64 ± 0.07 | 0.46 | 0.63 | 0.79 |
| | Specificity | 0.27 ± 0.09 | 0.04 | 0.29 | 0.67 |
| | F1 Score | 0.67 ± 0.05 | 0.55 | 0.67 | 0.75 |
| | Numbers of Predictors | 3 ± 1 | 2 | 2 | 5 |

| SVM Model | Performance Metrics | Summary Statistics |  |  |  |
| --- | --- | --- | --- | --- | --- |
|  |  | Mean ± Standard Deviation (SD) | Minimum | Median | Maximum |
| Modelling CRS Resilient and Vulnerability with FST | AUC | 0.53 ± 0.06 | 0.42 | 0.53 | 0.67 |
|  | Balanced Accuracy | 0.54 ± 0.05 | 0.43 | 0.53 | 0.62 |
|  | Sensitivity | 0.39 ± 0.10 | 0.16 | 0.39 | 0.63 |
|  | Specificity | 0.68 ± 0.06 | 0.54 | 0.67 | 0.80 |
|  | F1 Score | 0.36 ± 0.08 | 0.18 | 0.35 | 0.48 |
|  | Numbers of Predictors | 5 ± 1 | 3 | 5 | 6 |
| Modelling CRS Resilient and Vulnerability with EPM | AUC | 0.45 ± 0.06 | 0.30 | 0.44 | 0.56 |
|  | Balanced Accuracy | 0.46 ± 0.04 | 0.36 | 0.46 | 0.56 |
|  | Sensitivity | 0.64 ± 0.07 | 0.46 | 0.63 | 0.79 |
|  | Specificity | 0.27 ± 0.09 | 0.04 | 0.29 | 0.67 |
|  | F1 Score | 0.67 ± 0.05 | 0.55 | 0.67 | 0.75 |
|  | Numbers of Predictors | 3 ± 1 | 2 | 2 | 5 |

The repeated modelling approach selected an average of five predictors, with a maximum of six predictors. Figure 5A summarizes the frequency, importance and directionality of six predictors selected across 50 repetitions. All these connectivity features were selected more than 15 out of 50 runs of the algorithm. Two variables, including connectivity changes between hippocampal-cortical and corpus striatum-cortical network, and baseline connectivity between ventral striatum-cortical and corpus striatum-cortical network were selected in all 50 runs. The average importance of connectivity alteration between hippocampal-cortical and corpus striatum-cortical network ranked third, with a larger functional alteration indicating a higher probability of animals being vulnerable following CRS. By contrast, the average importance of baseline connectivity between ventral striatum-cortical and corpus striatum-cortical network ranked second, with a stronger baseline connectivity indicating a higher probability of animals being resilient following CRS. Each network connectivity feature only demonstrated a medium effect size between resilient and vulnerable groups *(Supplementary Figure S3)*. A substantial overlap of distributions in each predictor between these two groups was observed.

**Figure 5.**
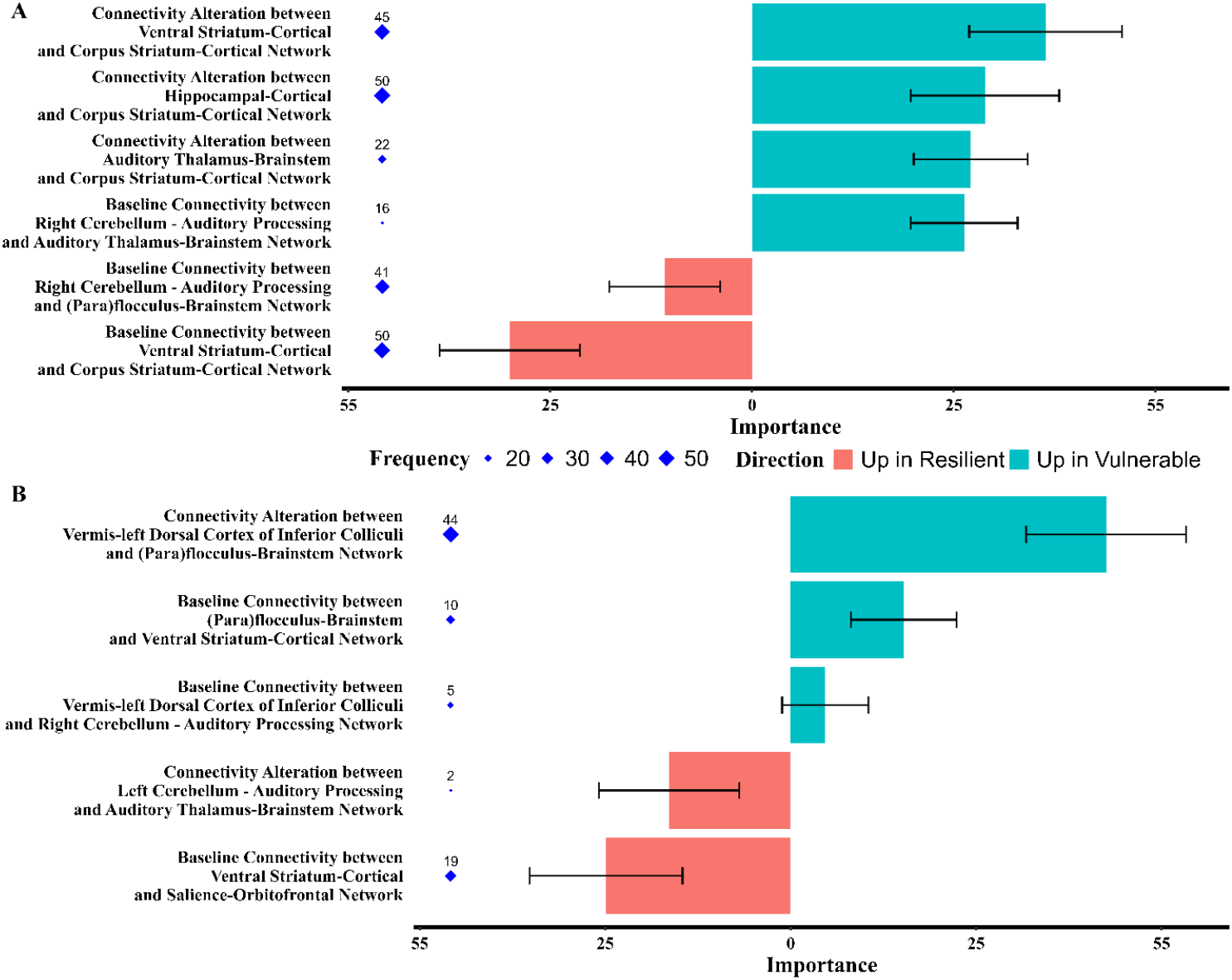
Variable Stability. **A). Neurobehavioral modelling of CRS resilience and vulnerability using FST:** divergent bar plot showing the importance (mean ± SD) and directionality of seven predictors. Different sizes of rhombus adjacent to the predictors showing the frequency of selection across 50 repetitions of nested cross-validation, with frequency labelled above rhombus; **B). Neurobehavioral modelling of CRS resilience and vulnerability using EPM:** divergent bar plot showing the importance (mean ± SD) and directionality of five predictors. Different sizes of rhombus adjacent to the predictors showing the frequency of selection across 50 repetitions of nested cross-validation, with frequency labelled above rhombus.

### Neurobehavioral Modelling of CRS Resilience and Vulnerability in EPM

Repeated nested cross-validation was conducted on 87 animals when modelling CRS resilience and vulnerability with EPM. The model also initially incorporated 156 independent variables, including baseline network connectivity and changes in the network connectivity. The model demonstrated poor performance to discriminate resilient animals from vulnerable animals with an average AUC of 0.45 and maximum AUC of 0.56 across 50 repetitions of nested cross-validation (*Table 1*). The F1 score achieved an average score of 0.67, ranging from 0.55 to 0.75. The balanced accuracy spanned from 0.36 to 0.56, with an average of 0.46. The average sensitivity of the model was 0.64 (range: 0.46 ~ 0.79), while the average specificity was 0.27 (range: 0.29 ~ 0.67).

The repeated modelling approach selected an average of three predictors, with a maximum of five predictors. None of these predictors were selected in all 50 runs (*Figure 5B*). But the connectivity alteration between vermis-left dorsal cortex of inferior colliculi and (para)flocculus-brainstem network were selected in 44 runs across 50 repetitions, with the highest importance. A bigger functional change suggested a higher probability of animals being vulnerable following CRS. Moreover, the average importance of baseline connectivity between ventral striatum-cortical and salience-orbitofrontal network ranked second, with a stronger baseline connectivity indicating a higher probability of animals being resilient following CRS. The effect sizes of these five predictors ranged from weak to medium between resilient and vulnerable groups *(Supplementary Figure S4)*. A substantial overlap of distributions in each connectivity feature between these two groups was observed.

## Discussion

The main aim of the present study was to identify connectome-based rs-fMRI biomarkers predictive of vulnerability and resilience following CRS. A key innovation of the study lies in the application of multivariate machine-learning approach to facilitate the identification of stable network connectivity features and patterns specific to each animal. The findings revealed no strong group effect sizes in behavioral changes following CRS within the same animals, suggesting the presence of resilient and vulnerable groups. Although the average model performance was close to chance level, baseline functional connectivity across cerebellum, brainstem, striatum, prefrontal and salience-orbitofrontal regions, as well as the functional alteration across hippocampus, striatum, prefrontal regions, auditory thalamus, cerebellum, inferior colliculi and brainstem were identified as stable features. These network connectivity features may provide insights into neuroimaging biomarkers to aid diagnosis and prevention of depression in humans.

### Behavioral Changes Following CRS

The lack of a strong group effect of CRS on FST and EPM behaviors indicates that physical restraint does not effectively induce stress and anxiety in all animals. The finding further supports the behavioral separation of resilient and vulnerable groups following chronic stress (2, 46). Interestingly, the observation that a higher proportion of animals were classified as CRS-vulnerable in the FST compared to the EPM can be explained by that the FST is a more stressful test. Moreover, the behavioral variability may reflect the high variability in the brain functional connectome reported in our previous publication (12). The individual variability in both behaviors and brain connectivity in the present and previous studies aligns with the literature showing significant variability in both animal models and humans with affective disorders (14, 47-49). Therefore, incorporating individual variation through multivariate machine-learning approach in the current work provides an opportunity to identify brain functional patterns that are relevant to understanding individual resilience and susceptibility to chronic stress.

### Modelling Performance

The poor modeling performance may indicate RSN connectivity could not provide accurate classification of CRS resilience and vulnerability using FST and EPM measures. However, the poor performance might be attributed to the class imbalance problem in small datasets because after applying SMOTE (the most popular oversampling method), the current models’ ability to identify the true minority class is still low (below 0.4; *Table 1*). This may further indicate that SMOTE’s effectiveness is limited by small sample sizes (50), or the complex resting-state functional connectome severely compromises the SMOTE’s ability to generate meaningful synthetic samples within the training process (51). Therefore, future studies with a larger sample size with balanced data are warranted to determine if network connectivity is useful to predict CRS resilience and vulnerability. Additionally, experiments incorporating optogenetics and MRI to capture the coactivation of brain regions during light stimulation and explore the role of specific circuits on behavior with machine learning will provide a better understanding of functional connectome in the rodent brain and may offer new insights into individual responses to stress.

### Features Predictive of Resilience and Vulnerability to CRS

Although the present prediction models struggled to achieve good performance, potentially because of insufficient and balanced data, several baseline connectivity and connectivity alterations following CRS were identified as stable features (*Figure 5*). The inclusion of baseline connectivity is pivotal as it has the potential to indicate an underlying resilience or susceptibility to developing passive coping and anxiety-related behaviors in response to stress. Moreover, the directionality of each selected feature seems to indicate the possibility of an animal developing resilience or susceptibility in response to CRS from a univariate perspective. However, the lack of strong group effect, along with substantial overlap in the data distribution of connectivity features between resilient and vulnerable groups highlights that the brain factors determining an animal’s response to a stressor and the impact of this response on behaviors are highly complex. The large overlap in the data distribution between resilient and vulnerable groups is also observed in studies investigating peak functional connectivity (14) and molecular markers (10). These findings further suggest that each feature alone could not differentiate the resilient from the vulnerable group. But complex changes across multiple functional regions as a composite pattern (52-54), could be useful to predict an individual’s response to chronic stress when the model performance is good. Importantly, the stability in feature selection across 50 repetitions of nested cross validation may indicate that the composite functional pattern consistently provides informative insights for predicting CRS resilience and vulnerability.

In the current work, the functional connectivity patterns linked to resilience and vulnerability to developing passive coping behavior following CRS differ from those associated with developing anxiety-related behavior. Baseline network connectivity across striatum, prefrontal regions, right cerebellum, auditory thalamus, (para)flocculus, and brainstem, and connectivity alteration across hippocampus, striatum, auditory thalamus, brainstem may serve as a composite pattern predictive of developing passive coping behavior. In contrast, baseline network connectivity across ventral striatum, cerebellum, inferior colliculi, salience-orbitofrontal regions and brainstem, along with connectivity alterations across cerebellum, inferior colliculi, auditory thalamus, and brainstem may act as a composite panel predictive of developing anxiety-related behavior.

It is challenging to compare the novel results with literature because this is the first study applying multivariate machine-learning approach to investigate stress resilience and vulnerability using FST and EPM measures. However, most of the functional regions identified in the present study have previously been implicated in stress responses in laboratory animals and in brain networks associated with depression (54-57). It is well-established that the hippocampal-cortical network is associated with dysfunctional cognitive and emotional processing in depression and other stress-related disorders. For example, a stronger connectivity between the hippocampus and ventromedial prefrontal cortex in humans is associated with a lower level of stress resilience during Covid-19 pandemic (58). Moreover, the prefrontal cortex, hippocampus and ventral striatum are key regions of a dynamic network underlying vulnerability to chronic stress in mice (54). Although cerebellum is traditionally associated with motor coordination and learning, increasing evidence has shown that cerebellum-brainstem (particularly ventral tegmental area) network contributes to developing depression-like behaviors induced by chronic stress in rodents (57). As anxiety and passive coping behaviors are highly co-occurrent with depression (59), the functional pattern identified here may further shed light on neuroimaging biomarkers to aid diagnosis and prevention of depression in humans. Finally, it is worth reiterating that the results should be interpreted with caution given that the performance of the current model is close to chance level and further validation in a larger and balanced dataset is warranted.

### Limitations and Future Directions

Several limitations are worth mentioning. First, the data was collected over a period of two years in several cohorts. Although the experimental protocols and housing conditions were consistent, variation could arise from changes in student experimenters conducting the procedures and uncontrollable factors at the level of the commercial suppliers, such as breeding conditions, parental care, and genetic drift. Second, in an effort to reduce variability, only male rats were used in the study. However, this decision unfortunately contributes to the significant bias against using female animals in medical research. It will be important to include female rodents in future multivariate studies because women are twice as likely as men to develop anxiety (60). Moreover, animals started the CRS intervention during emerging adulthood, a period when the brain is still undergoing structural and functional changes following adolescence. Therefore, there could be age-dependent effects in the present study, although two of our previous publications have shown that behaviors (20) and network connectivity (12) did not change significantly over time in a control group of animals that did not undergo CRS. Nevertheless, the aim of this study was to compare vulnerable and susceptible animals within an intervention, a design that does not require a separate control group. Finally, the translational relevance of behavioral tests in animal models of anxiety, stress and depression remains a key limitation in the field. The current findings need to be interpreted with caution and replicated in other models.

## Conclusion

The present study is the first to identify rs-fMRI biomarkers potentially predictive of stress resilience and vulnerability to developing anxiety-related and passive coping behaviors in an animal model of depression. The current findings highlight the benefits of shifting the focus of biomarker research from the group to the individual level (61, 62). Although the biomarkers identified here have poor predictive validity, the stability of several connectivity features could support the advancement of personalized prevention, diagnosis, and treatment interventions for depression. Additionally, the success of brain-behavior modeling and translational neuroscience research depends on clinical relevance and accurate measurements of captured behaviors. Therefore, refining methodology to objectively measure clinically relevant behaviors and emotions in animals is a priority in neuroscience research.

## Supporting information

Supplementary Text and Scripts

Supplementary Workspace

## Acknowledgments

JR is supported by Perron Institute for Neurological and Translational Science. TD is supported by the Australian Government International Research Training Program scholarship, and Byron Kakulas Prestige scholarship.

TD and JR conceived and designed the study. TD processed and analyzed all the data, as well as wrote the first draft of the manuscript. SDA, MS and AZ provided advice and input regarding brain-behavior modelling. All authors reviewed the article and approved the submitted version.

