## Supplementary Text and Scripts for "Identifying Neuroimaging Biomarkers of Resilience and Vulnerability to Chronic Stress in An Animal Model: An Exploratory Analysis"

Dai *et al.*

### Supplementary Text

#### Methods and Materials

##### *Experimental Procedures*

###### Chronic Restraint Stress

CRS procedure adhered to the protocol specified in Ulloa *et al.* (1). Detailed description of the protocol for the present study can be found in Dai *et al.* (2) and Seewoo *et al.* (3). Briefly, each rat was placed in a transparent acrylic tube to restrict their free movement for 2.5 h daily, for 13 consecutive days. Following CRS, rats were returned to their home cages.

###### Elevated Plus Maze Test

EPM test was conducted following the protocols described in Walf and Frye (4) and Seewoo *et al.* (3) to evaluate anxiety-related behaviors. The EPM apparatus was plus-shaped consisting of two open and two closed arms, and a center zone (10 × 10 cm). Each arm has a length of 50 cm and width of 10 cm. Animals were positioned initially in the center zone facing the open arm and then allowed to freely explore the maze for five minutes. One animal was tested at a time. The apparatus was wiped with 70% ethanol to remove olfactory cues after each test session.

###### Forced Swimming Test

FST was carried out as detailed in Slattery and Cryan (5) and Seewoo *et al.* (3) to evaluate learned-helplessness – one feature of depression-related behaviors. Animals were individually placed in a 20-Liter opaque white bucket (41 cm height × 28 cm width) filled with water (30 cm depth, 23 - 25°C), for six minutes. The bucket was emptied, rinsed, and wiped with 70% ethanol to remove olfactory cues after each test session.

###### MRI Acquisition

The animal anesthesia and MRI acquisition protocols were described and discussed thoroughly in the previous publications (2). Briefly, rats were anesthetized under combined use of isoflurane and medetomidine. Anesthesia was inducted with 4% isoflurane and reduced to 2% once the respiratory rate dropped to 55 - 60 breaths/min. Medetomidine was administered subcutaneously, with an initial bolus injection (0.05 - 0.1 mg/kg) and continuous infusion at 0.15 mg/kg/hr after animals were stabilized on 2% isoflurane for two minutes. Meanwhile, isoflurane was slowly reduced to 0.5% - 0.75% based on animal's respiratory rate. The average time elapsed following the induction of medetomidine was 32 ± 4 minutes. Under anesthesia, rats were scanned using a 9.4T Bruker Biospec 94/30 pre-clinical MRI scanner (Bruker BioSpin GmbH, Germany). The raw images of structural MRI and rs-fMRI, each with 21 coronal slices, were compiled in one data package.

The structural MRI was acquired using an accelerated multi-slice 2D rapid acquisition with relaxation enhancement sequence (thickness = 1 mm, matrix = 280× 280, voxel size = 0.1× 0.1 $mm^2$, acceleration = 8, TR = 2500 ms, TE = 33 ms). The rs-fMRI was acquired using a single-shot gradient-echo echoplanar imaging (EPI) method (thickness = 1 mm, 300 volumes, matrix = 94 × 70, pixel size = 0.3 × 0.3 $mm^2$, TR = 1500 ms, TE = 11 ms, order automatic ghost correction = 1, receiver bandwidth = 300 kHz).

##### *Data Processing*

###### FST Manual Annotation

FST videos were blinded and manually annotated by researchers. Each FST recording was first renamed with a unique random number and then manually annotated using a time-sampling approach (5). The first five minutes of each recording was split into 60 five-second clippings, and the predominant behavior in each clipping was rated. Four behaviors were scored: a). Immobility/Floating: keeping their head above water with minimal movements and without struggling; b). Swimming: performing horizontal movements, crossing over into another quadrant and diving; c). Climbing: finding supports along the bucket wall with forepaws moving upwards; d). Latency: the time (in second) elapsed until animals exhibit their first floating behavior. If animals were floating horizontally or managed to escape more than once during their test session, they were excluded from the analysis. Only 79 animals (46 from Hennessy et al. (14) and Seewoo et al. (15), and 33 unpublished) completed both FST sessions, with a total of 158 recordings annotated. Raw values of these four behaviors at baseline and post-CRS for 79 animals can be found in Supplementary Spreadsheet.

##### MRI Batch Processing

###### *A). Pre-Processing*

The pre-processing for each data package was batch processed as follows: a). Organize data to the Brain Imaging Data Structure (BIDS; 6) folder structure using BrkRaw (version: 0.3.5; 7); b). Reorient raw images in the radiological (left-anterior-superior) view; c). Perform bias field correction for anatomical images with 3D Slicer (version: 4.8.1; 8); d). Extract the brain for anatomical images and prepare an individualized brain mask for the next step; e). Strip the skull for functional images using the brain mask; f). Upscale the voxel size of functional images by a factor of 10, with a voxel size of 3 × 3 × 10 mm$^3$ (9, 10); g). Inspect the quality of reorientation and brain extraction visually with FSL/slices (no images failed to reorient and extract brain within the batch process); h). Denoise upscaled functional brain images using FSL/MELODIC (version: 3.15; 11) and FMRIB's ICA-based Xnoiseifier (FIX; version: 1.068; 12, 13). This step is described thoroughly in method 2.3.2 in the previous work (2); i). Each de-noised functional brain image was registered to a Sprague-Dawley rat brain atlas, which was down sampled by a factor of eight (9, 10) from the Waxholm Space atlas (RRID:SCR_017124; 14), to construct de-noised and normalized functional images. The down-sampled atlas has a voxel size of 3.125 × 3.125 × 3.125 mm$^3$ and all subsequent processing was performed in this atlas space.

###### *B). Optimal and Reproducible Functional Component Identification*

In addition to the 96 animals mentioned in the study design, 13 more animals underwent MRI sessions at baseline without any intervention throughout the timeframe of the experiment. These 13 animals' baseline images (also de-noised and normalized) were included in this step to generate a more representative brain template (15) for functional component identification. So, a total of 109 animals were used to identify the optimal number of functional components of the rat brain. Moreover, the rationale for only including baseline data is that CRS-related resting-state alterations compromise the identification of functional component template (10).

The group independent component analysis (group-ICA) combining temporal concatenation approach (16) in the mICA toolbox (17) was applied to identify the optimal number of functional components for the whole brain gray matter of the rats. The rationales of performing this analysis have been discussed thoroughly in the previous work (2). Briefly, all de-noised and normalized functional images at baseline (N = 109) and the atlas mask of whole brain gray matter were imported to the mICA toolbox. Random split-half sampling was performed at 30 different levels of dimensionality ranging from 10 to 300 components with an interval of 10, with 50 repetitions. For each repetition, MELODIC group-ICA was first conducted on both split-half groups (N = 54 per group) to identify functional components, respectively. Then Pearson's correlation was used to generate a cross-correlation matrix between intergroup components, which were later matched through maximizing the summed correlation of all component pairs with Hungarian sorting algorithm (18). Mean correlation with 95% confidence interval (CI) were calculated over 50 repetitions. This analysis was repeated for three different Gaussian kernels of full-width half maximum (FWHM) at threefold, fourfold, fivefold voxel size (corresponding to 9.375, 12.5, and 15.625 mm) of the down-sampled atlas (19, 20). Resultant correlation outputs of 90 combinations (30 levels of dimensionality with three Gaussian kernels) were recorded in the *Supplementary Spreadsheet* and presented in a curve plot. The optimal group-ICA dimensionality under three Gaussian kernels was determined based on the global maximum of correlation outputs (17). This means that a combination with the maximum correlation value presented in the curve plot was considered as an optimal dimensionality.

Because mICA toolbox cannot differentiate true reproducible components from noise-related components, group-ICA outputs of all split-half sampling groups (N =100) at the resultant dimensionality and Gaussian kernel were fed to the gRAICAR toolbox (21) in MATLAB (version: r2019b) to identify true reproducible functional components. Detailed explanation can be found in method 2.3.3.3 in the previous publication (2). Briefly, the toolbox first ranked and aligned functional components according to their reproducibility over repeated ICA realizations. For each aligned and ranked component/node, if the ratio of significant groups was greater than or equal to 0.01, an averaged spatial map over multiple realizations was then generated. Spatial maps of these nodes can be found in *Supplementary Spreadsheet*. Explanation of the significance of a group and ratio of significant groups see Yang *et al.* (21). A higher ratio of significant groups suggests a stronger inter-group consistency. Noise-related nodes (the ratio of significant groups of that node was lower than 0.01) were excluded from the next processing.

Spatial maps of reproducible functional components/nodes were then concatenated using FSL/fslmerge to construct a 4D image. Rat brain parcellation was conducted by applying mixture modelling approach (FSL/melodic; 22) and Ward's algorithm (Nilearn; version: 0.10.1; 23, 24) to the 4D image. As a result, the spatial boundary of each functional node was determined. A 2D image of each node was generated using FSL/slices_summary. All the nodes were finally merged to create a group-ICA template of the whole brain gray matter and ready for the next processing.

#### Results

##### *Optimal and Reproducible Functional Component*

As shown in Supplementary Figure S1A, the correlation value (0.78 ± 0.01) at the dimensionality of 220 and Gaussian kernels of FWHM 15.625 mm was highest. As such, 220 was considered the optimal dimensionality for further analysis to identify reproducible functional components. Additionally, the dimensionality-correlation curve revealed that higher spatial smoothing led to a higher correlation of group-ICA spatial maps at each dimensionality from 10 to 300. Under three Gaussian kernels, the correlation value initially dropped and then increased from the dimensionality of 30.

The ratio of significant groups for 220 independent components were then evaluated using gRAICAR over 100 groups. Supplementary Figure S1B showed that all 100 groups contributed to 190 components, with no group contributing to three components. As such these three functional components were noise-related components. Moreover, 217 nodes had a ratio of significant groups greater than or equal to 0.01, and thus were considered as true and reproducible functional components for brain parcellation and clustering analysis.

##### *Hierarchical Clustering*

The data-driven approach - hierarchical clustering merged 217 nodes into 13 large-scale functional networks in the rat brain (*Supplementary Figure S2 & Figure 4*). These networks included salience-orbitofrontal, lateral-temporal cortical, hippocampal-cortical, default mode-somatomotor, spatial attention, corpus striatum-cortical, ventral striatum-cortical, hypothalamus-thalamus, auditory thalamus, (para)flocculus-brainstem and three cerebellum-auditory processing networks. Spatial symmetry was presented in some homologous brain regions within the majority of these networks (detailed description and discussion can be found in the *Supplementary Text*).

**Salience-Orbitofrontal Network (11 nodes)**. The network was mostly cortical, except that bilateral anterior claustrum and a proportion of right nucleus accumbens were involved. Two core salience-related nodes – anterior ventral insula (dorsal and ventral area of the agranular insula) and anterior cingulate cortex presented asymmetrical resting-state activity. The anterior insula had larger spatial extent of resting-state activity in the left than right hemisphere, while the anterior cingulate exhibit oppositely. Similarly, the spatial extent of resting-state activity in the orbitofrontal area was broader in the right than left hemisphere. Overall, the right hemisphere of this network had a broader spatial extent of resting-state activity than that of the left.

**Lateral-Temporal Cortical Network (8 nodes)**. The network was completely cortical, including entorhinal, auditory, posterior and parietal parts of dysgranular and granular insula, perirhinal, and temporal association cortex. There was an overall larger spatial extent of resting-state activity in the right than left lateral-temporal hemisphere.

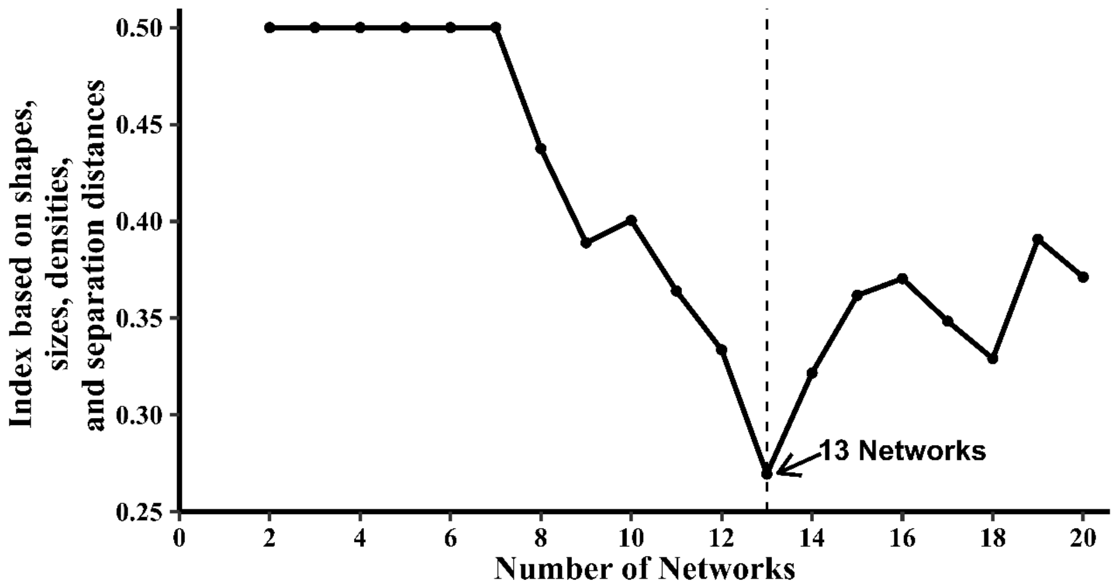


**Supplementary Figure S2. Identification of optimal numbers of large-scale functional networks in the rat brain.** The lowest SSDS index was 0.27, indicating that 13 networks were the optimal number and natural division for 217 functional nodes.

**Hippocampal-Cortical Network (12 nodes).** The network comprised ventral hippocampus and its adjacent brain regions. The resting-state activity of the ventral hippocampus was mostly symmetrical. Although the entorhinal cortex and amygdala demonstrated broader spatial extent in the left than right hemisphere, the perirhinal cortex presented oppositely and the postrhinal cortex only presented unilateral resting-state activity in the left hemisphere.

**Default Mode-Somatomotor Network (27 nodes)**. The network was completely cortical. Two important default mode components – the posterior cingulate and anterior retrosplenial cortex demonstrated bilateral symmetry in the resting-state activity. But the resting-state activity of prelimbic cortex was completely absent in the right hemisphere. Moreover, the spatial extent of resting-state activity in the visual and posterior area of parietal association cortex was larger in the left hemisphere, whereas sensorimotor structures were mostly symmetrical.

**Spatial Attention Network (21 nodes)**. This network consisted of both subcortical and cortical components. The superior colliculi and dorsal hippocampus demonstrated symmetrical resting-state activity, whereas inferior colliculi and pulvinar had broader spatial extent of resting-state activity in the left than right hemisphere. In contrast, cortical areas, including posterior retrosplenial, visual and auditory cortex displayed larger spatial extent in the right than left hemisphere.

**Corpus Striatum-Cortical Network (23 nodes)**. The network included caudate putamen, globus pallidus, and adjacent cortical areas. The resting-state activity of the caudate putamen and somatomotor cortices were mostly symmetrical. By contrast, the spatial extent of resting-state activity in the globus pallidus, insula and orbitofrontal cortex was broader in the left than right hemisphere. Additionally, the amygdala and piriform cortex in this network only demonstrated unilateral resting-state activity in the left hemisphere.

**Ventral Striatum-Cortical Network (24 nodes)**. The network consisted of nucleus accumbens, ventral pallidum, basal forebrain, and adjacent cortex. Nucleus accumbens demonstrated larger spatial extent in the left than right hemisphere, whereas the ventral pallidum, ventral striatal region, basal forebrain and piriform cortex presented oppositely.

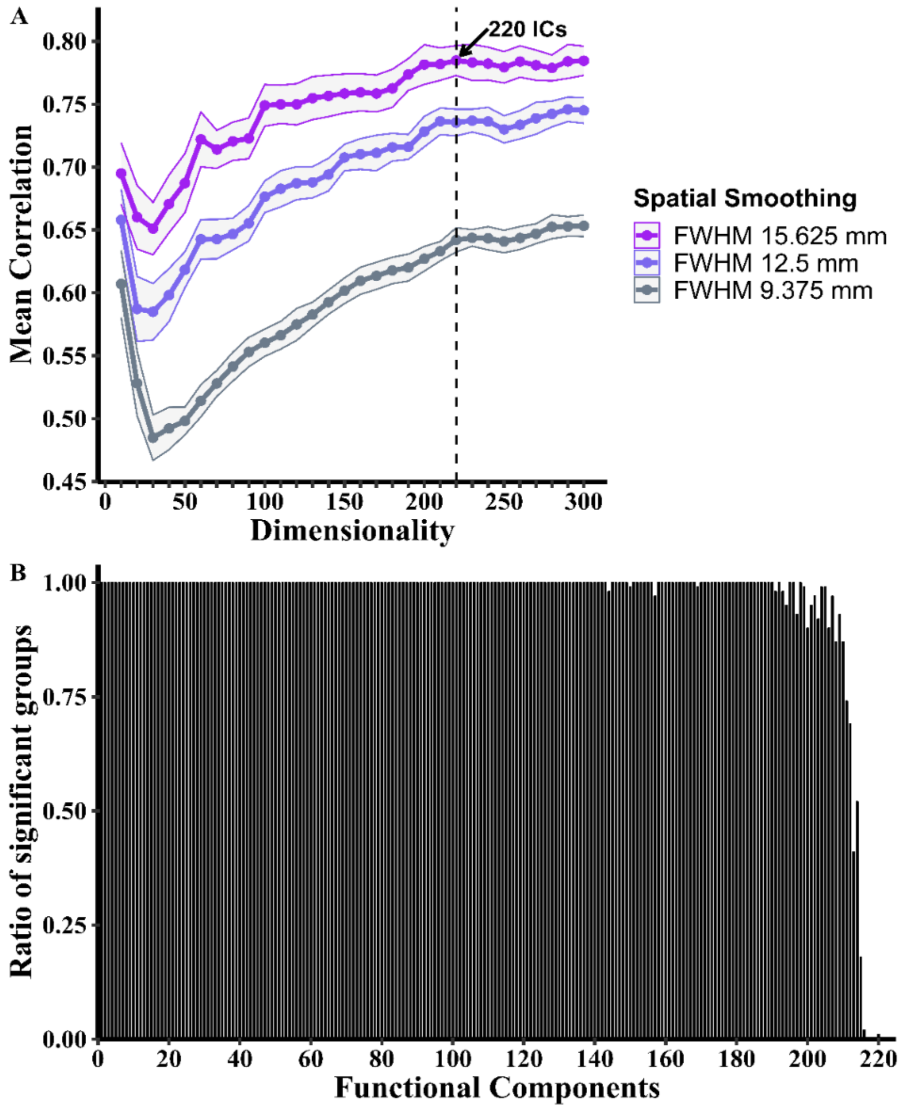


**Supplementary Figure S1. Optimal and reproducible functional components in the rat brain. A)**. Curve plot showing the spatial smoothing effects on determining the optimal number of functional components. The dimensionality-correlation curves under three different Gaussian kernels showed that an increase in the spatial smoothing resulted in an overall increase in the correlation of group-ICA spatial maps. The optimal number of components was 220 independent components (ICs; arrow), which was the global maximum derived from the curves. The Curves were plotted with mean ± 95% CI; **B)**. Column chart showing the proportion of groups contributed to each aligned component at the dimensionality of 220 following RAICAR over 100 group-ICA spatial maps. 217 nodes had a ratio of significant groups greater than or equal to 0.01, and thus were considered as reproducible functional components.

**Hypothalamus-Thalamus Network (11 nodes)**. The network was subcortical, including hypothalamus and dorsal thalamus, such as intermediodorsal thalamic and posterior thalamic nucleus. Their resting-state activity was mostly symmetrical.

**Auditory Thalamus-Brainstem Network (16 nodes)**. The network mainly contained the medial geniculate body and brainstem. The resting-state activity differed depending on the division of the medial geniculate body, with dorsal division showing broader spatial extent in the right than left hemisphere, medial and suprageniculate division presenting larger spatial extent in the left. By contrast, the resting-state activity in the ventral division and marginal zone of the medial geniculate body, along with brainstem was mostly symmetrical.

**(Para)flocculus-Brainstem Network (24 nodes)**. The resting-state activity of the para-flocculus, flocculus, and brainstem involved in this network was mostly symmetrical.

**Three Cerebellum-Auditory Processing Networks**. In the vermis-left dorsal cortex of inferior colliculi (16 nodes), the vermis had larger spatial extent of resting-state activity in the left than right hemisphere, with dorsal cortex of inferior colliculi displaying unilateral resting-state activity in the left hemisphere. As for the right cerebellum - auditory processing network (16 nodes), the spatial extent of resting-state activity in the cerebellum and inferior colliculi was broader in the right than left hemisphere. In contrast, in the left cerebellum - auditory processing network (8 nodes): the resting-state activity of cerebellum and inferior colliculi was completely absent in the right hemisphere.

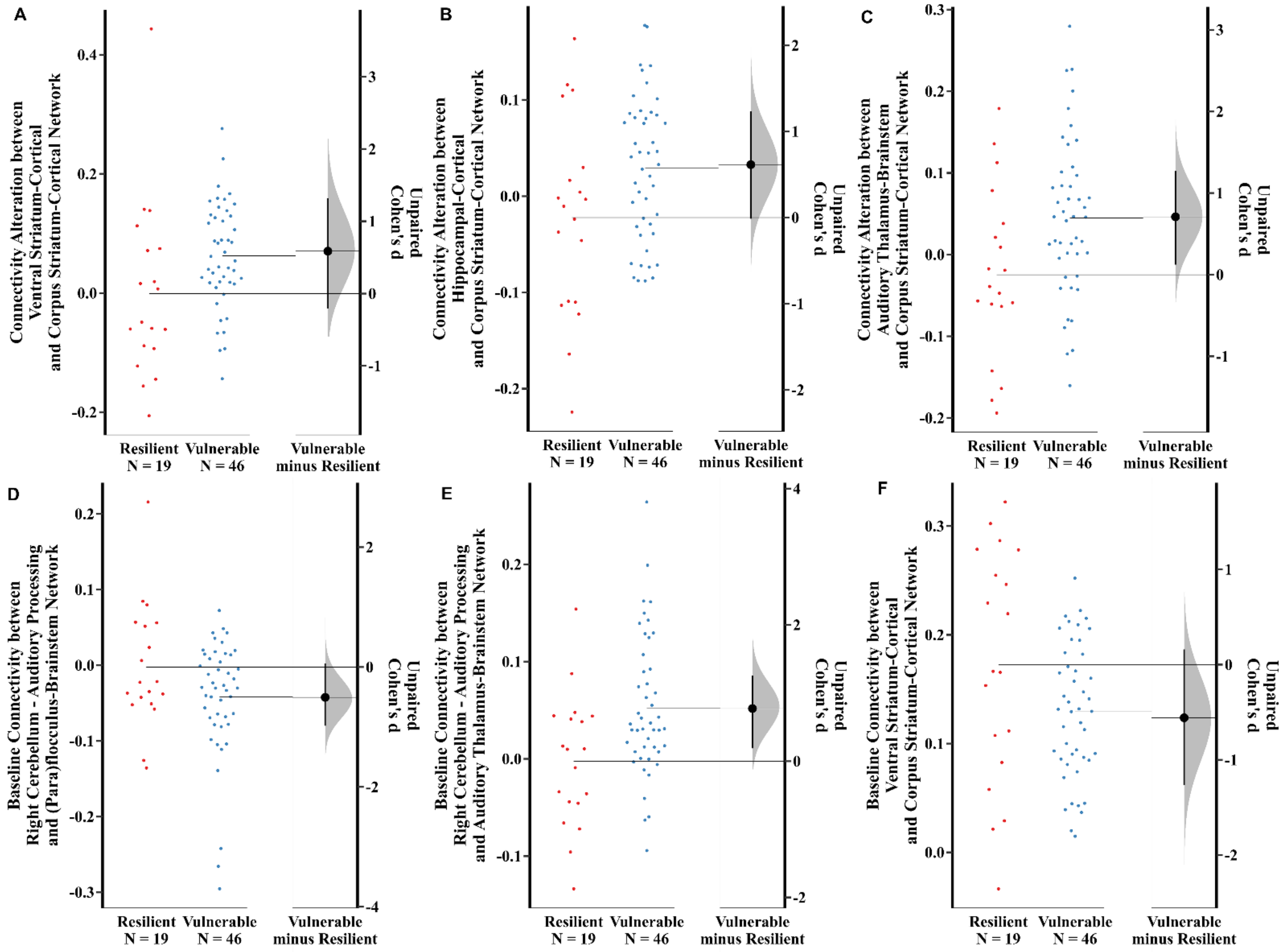


**Supplementary Figure S3. Group Effect Size of Seven Predictors in Neurobehavioral Modelling of CRS Resilience and Vulnerability Using FST.**

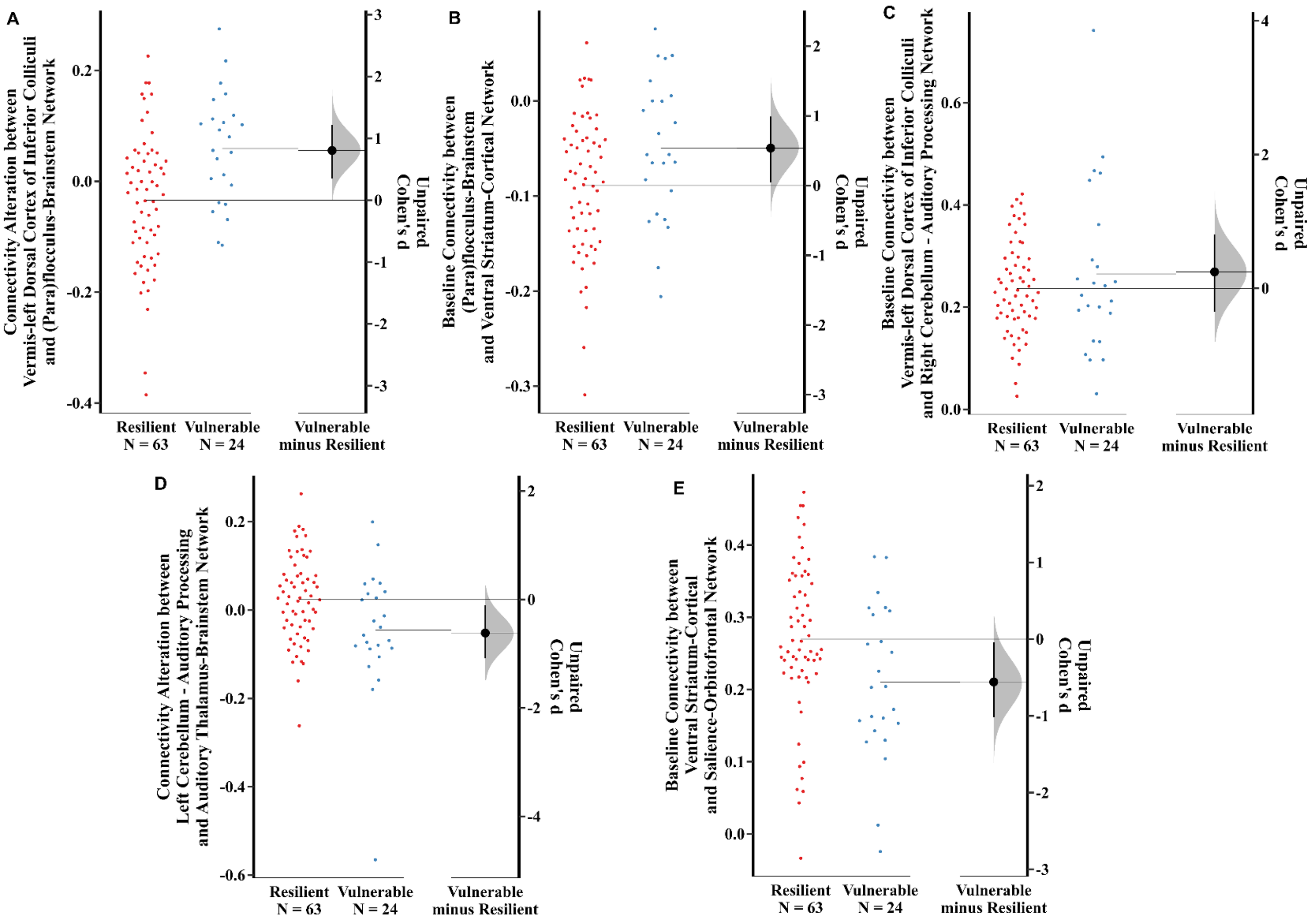


**Supplementary Figure S4. Group Effect Size of Five Predictors in Neurobehavioral Modelling of CRS Resilience and Vulnerability Using EPM.**

#### Discussion

##### *Large-scale Resting-state Functional Networks*

The 13 large-scale functional networks classified in the present study show some similarity to the functional organizations observed in healthy humans and rodents in the literature (29, 34-38). Similarities and differences of functional organization are discussed below.

**Salience-Orbitofrontal Network.** The anterior ventral insula and anterior cingulate cortex in this network are core hubs of salience network observed in humans (39) and rodents (38). The involvement of orbitofrontal cortex can be explained by its anatomical connections with insular and cingulate cortex (40), functional overlaps between orbitofrontal and insular cortex (41), and spatial closeness. Both salience network and orbitofrontal cortex are believed to participate in processing affective salience to regulate emotion (40, 42). The broader resting-state activity in the right hemisphere of this network agrees with the right-hemispheric dominance hypothesis of affective process and emotion in humans (43, 44).

**Lateral-Temporal Cortical Network.** The network has not been reported previously, but this functional formation can be explained by the anatomical and functional connections among involved cortices. The parietal insula receives projections from auditory cortices (45). Both posterior and parietal insula send substantial inputs to the para-hippocampal regions. Moreover, the parietal insular, perirhinal and temporal association cortex are reported to engage in the auditory fear-conditioning to both intermittent and continuous tones (45-47).

**Hippocampal-Cortical Network.** The network has not been observed in other rs-fMRI studies, but this functional organization is supported by anatomical connections among involved subcortical and cortical structures, as well as their functions. The ventral hippocampus has complex and extensive interconnections with amygdala and para-hippocampal regions (48-50). These interconnections are believed to play a key role in modulating affects, emotions, stress, and emotional memory storage.

**Default Mode-Somatomotor Network.** The core DMN structures (posterior cingulate and anterior retrosplenial cortex) identified here are consistent with previous publication (2) based on a hierarchical analysis. The result also agrees with other rodent studies using group ICA analysis with a low dimensionality from 10 to 30 or seed-based approaches (37, 51-57). The somatosensory structures being dependent from the DMN observed here is similar to the previous and other rodent studies (2, 52, 53, 57), but not in Zerbi *et al.* (37). However, the integration of somatomotor areas in the DMN is not surprising because DMN coordinates with multiple brain regions for passive sensory processing (58). The involvement of hippocampal regions in the DMN has been controversial. The present study has shown that hippocampus is absent from the DMN, which is similar to some studies (51, 55, 56), but different from others (2, 37, 52, 54).

**Spatial Attention Network.** The components of this network are similar to what has been observed in awake rodents (29, 59). The superior colliculi and pulvinar have been well-known to be involved in spatial visual attention (60). Dorsal hippocampus sends direct projections to retrosplenial cortex, contributing to spatial memory (61, 62). Additionally, the asymmetrical resting-state activity of the retrosplenial, visual, and auditory cortex in this network is supported by the right-hemispheric dominance of spatial attention in humans (63, 64).

**Corpus Striatum-Cortical Network and Ventral Striatum-Cortical Network.** These two networks belong to basal ganglia networks. However, the integration of cortical areas with neither corpus striatum nor ventral striatum is observed in previous rodent and human studies (29, 35, 65). The striatum, insula, orbital cortex, and amygdala involved in both networks form cortico-amygdala-striatal circuit. The circuit mediates emotions and affects through collectively processing attentional states and salient stimuli in both rodents and primates (66, 67). Unfortunately, it is difficult to explain the resting-state separation of the cortico-amygdala-striatal circuit in these two networks.

**Hypothalamus-Thalamus and Auditory Thalamus-Brainstem Network.** These two networks differ from other rodent studies (9, 29, 53) showing hypothalamus and thalamus as two independent functional properties. However, it is more biologically reasonable to see the coactivation of hypothalamus and thalamus in one network, as well as medial geniculate body and brainstem in the latter. This is because it's unlikely that hypothalamus and thalamus functionally act on their own, without the involvement of other brain regions. Hypothalamus is well known in its role of stress regulation and autonomic control (68). Several thalamic nuclei, such as the paraventricular and mediodorsal nuclei, are also involved in autonomic control. This may explain the functional integration of hypothalamus and thalamus observed in this study. As for the auditory thalamus-brainstem network, medial geniculate body (core component) is well characterized as major synaptic centres receiving massive input from auditory stations of midbrain - the inferior colliculi and nuclei of the lateral lemniscus (69).

**(Para)flocculus-Brainstem Network and Three Cerebellum-Auditory Processing Networks.** Cerebellum splitting into four Networks in the current work is different from one study showing one large independent component of cerebellum resulted from a group ICA analysis with a low dimensionality of 20 (29). The present result is also not in line with another study reporting two cerebellar networks by apply a hierarchical clustering analysis with 122 pre-selected brain regions (37). However, the separation of cerebellum observed here is supported by a human study demonstrating the distinctness of cerebellar contributions to other functional organizations in the brain(70). Considering involved brain structures, these four networks seem to implicate their role in auditory processing (71).

A total of five networks, including auditory thalamus-brainstem, (para)flocculus-brainstem, and three cerebellum-auditory processing networks, are related to auditory processing in non-cortical areas. Studies have shown that neurophysiological recording can still detect neural activities in the auditory midbrain and cortex in response to tones while rodents are anaesthetized (72, 73). These suggest that anesthetized animals can still hear the noise from MRI machine during their scan, resulting in the substantial activation of brain regions involved in auditory processing. In addition, the inconsistencies of abovementioned functional networks comparing to the literature can be attributed to the methodological differences in detecting large-scale functional networks.

### Supplementary Script

#### MRI Batch Processing

#### *A). Pre-Processing*

##### a). Organize data to the BIDS structure

```
#!/bin/sh
## each raw package was downloaded and compressed in tar file, with the same naming pattern of
"AnimalID_Timepoint". The command is to unzip the tar file and rename the PvDatasets and metadata
files.
for folder in PATH/TO/Cohort*/*; do cd $folder && echo $folder && for file in *.tar; do echo $file &&
foldername=${file%".tar"} && mkdir $foldername && tar -xvf $file -C $foldername && mv
$foldername/*/*/*PvDatasets $foldername/*/*/metadata.txt $foldername && rm -r $foldername/000* &&
mv $foldername/*PvDatasets $foldername/$foldername.PvDatasets && mv $foldername/metadata.txt
$foldername/"$foldername"_metadata.txt;done;done
## using Brkraw to create BIDS structures for each animal in a virtual python environment for each
cohort separately
source PATH/TO/Twainpythonenv/bin/activate
cd PATH/TO/Cohort1
for folder in *; do brkraw bids_helper $folder "$folder"_bids_datasets -j; done
brkraw bids_convert Cohort1 Cohort1/Cohort1_bids_datasets.csv -j Cohort1/Cohort1_bids_dataset.json
-o Bids_folder
```

##### b). Reorient raw images in the radiological

```
#!/bin/sh
## Reorient T2w
module load fsl/6.0.3
for folder in PATH/TO/Bids_folder/sub-Cohort*/ses*/anat; do
ID=${folder#*Bids_folder/}
ID=${ID%/anat}
ID=${ID////_}
echo $ID
cp "$ID"_acq-coronal_T2w.nii.gz "$ID"_T2w_coronal.nii.gz
fslswapdim "$ID"_T2w_coronal.nii.gz -x z y "$ID"_T2w_coronal.nii.gz
fslorient -copysform2qform "$ID"_T2w_coronal.nii.gz
cp "$ID"_T2w_coronal.nii.gz "$ID"_T2w_coronal_x10.nii.gz
fslchpixdim "$ID"_T2w_coronal_x10.nii.gz 1 10.5 1
## Reorient rs-fMRI data
for folder in PATH/TO/Bids_folder/sub-Cohort*/ses*/func; do
   [ -d "$folder" ] && cd "$folder" &&
#get animal ID
ID=${folder#*Bids_folder/}
ID=${ID%/func}
ID=${ID////_}
echo $ID
cp "$ID"_acq-bold_EPI.nii.gz "$ID"_rs_bold.nii.gz
fslswapdim "$ID"_rs_bold.nii.gz -x z y "$ID"_rs_bold.nii.gz
fslorient -copysform2qform "$ID"_rs_bold.nii.gz
done
```

**c). Perform bias field correction for anatomical images with 3D Slicer**

```
#!/bin/sh
for folder in PATH/TO/Bids_folder/sub-Cohort*/ses*/anat; do
        [ -d "$folder" ] && cd "$folder" &&
ID=${folder#*Bids_folder/}
ID=${ID%/anat}
ID=${ID////_}
echo $ID
/usr/local/3dslicer/4.8.1/Slicer --launch /usr/local/3dslicer/4.8.1/lib/Slicer-4.8/cli-
modules/N4ITKBiasFieldCorrection --meshresolution 1,1,1 --splinedistance 0 --bffwhm 0 --iterations
50,40,30 --convergencethreshold 0.0001 --bsplineorder 3 --shrinkfactor 4 --wienerfilternoise 0 --
nhistogrambins 0 "$ID"_T2w_coronal.nii.gz "$ID"_T2w_coronal_BFC.nii.gz
```

**d). Extract the brain for anatomical images and prepare an individualised brain mask**

```
#!/bin/sh
module load fsl/6.0.3
for folder in PATH/TO/Bids_folder/sub-Cohort*/ses*/anat; do
        [ -d "$folder" ] && cd "$folder" &&
ID=${folder#*Bids_folder/}
ID=${ID%/anat}
ID=${ID////_}
echo $ID
if test -f "$ID"_T2w_coronal_brain.nii.gz; then
   echo ""$ID"_T2w_coronal_brain.nii.gz exist"
else
       echo ""$ID"_T2w_coronal_brain.nii.gz does not exist"

echo doing corregistrations
flirt -in "$ID"_T2w_coronal_BFC.nii.gz -ref PATH/TO/Atlas/atlas_brain.nii.gz -out
"$ID"_T2w_coronal_hratlas_brain.nii.gz -omat "$ID"_T2w_coronal_hratlas_brain.mat -bins 30 -cost
corratio -searchrx -90 90 -searchry -90 90 -searchrz -90 90 -dof 9  -interp trilinear
#get inverse matrix
convert_xfm -omat "$ID"_T2w_coronal_hratlas_brain_inverse.mat -inverse
"$ID"_T2w_coronal_hratlas_brain.mat
#get brain mask in t2 space and extract t2
flirt -in PATH/TO/Atlas/atlas_brain.nii.gz -ref "$ID"_T2w_coronal_BFC.nii.gz -out
"$ID"_T2w_coronal_hratlas_brain_inverse.nii.gz -init "$ID"_T2w_coronal_hratlas_brain_inverse.mat -
applyxfm
fslmaths "$ID"_T2w_coronal_hratlas_brain_inverse.nii.gz -bin
"$ID"_T2w_coronal_hratlas_brain_inverse_mask.nii.gz
fslmaths "$ID"_T2w_coronal_BFC.nii.gz -mas "$ID"_T2w_coronal_hratlas_brain_inverse_mask.nii.gz
"$ID"_T2w_coronal_brain.nii.gz
rm "$ID"_T2w_coronal_hratlas_brain.nii.gz
fslmaths "$ID"_T2w_coronal_brain.nii.gz -bin "$ID"_T2w_coronal_brain_mask.nii.gz
fi
done
```

**e&f). Strip the skull for functional images**

```
#!/bin/sh
module load fsl/6.0.3
echo doing bet
for folder in PATH/TO/Bids_folder/sub-Cohort1*/ses*/func; do
        [ -d "$folder" ] && cd "$folder" &&
ID=${folder#*Bids_folder/}
ID=${ID%/func}
ID=${ID////_}
echo $ID
if test -f "$ID"_rs_bold_brain_x10.nii.gz; then
   echo ""$ID"_rs_bold_brain_x10.nii.gz exist"
else
       echo ""$ID"_rs_bold_brain_x10.nii.gz does not exist"
echo "extract rs"
flirt -in "$ID"_rs_bold.nii.gz -ref ../anat/"$ID"_T2w_coronal.nii.gz -out "$ID"_rs_t2.nii.gz -omat
"$ID"_rs_t2.mat -bins 30 -cost corratio -searchrx 0 0 -searchry 0 0 -searchrz 0 0 -dof 6 -interp trilinear
fslmaths "$ID"_rs_t2.nii.gz -mas ../anat/"$ID"_T2w_coronal_brain_mask "$ID"_rs_t2_brain.nii.gz
convert_xfm -omat "$ID"_rs_t2_inverse.mat -inverse "$ID"_rs_t2.mat
flirt -in "$ID"_rs_t2_brain.nii.gz -ref "$ID"_rs_bold.nii.gz -out "$ID"_rs_t2_brain_inverse.nii.gz -init
"$ID"_rs_t2_inverse.mat -applyxfm
fslmaths "$ID"_rs_t2_brain_inverse.nii.gz -bin "$ID"_rs_mask.nii.gz
fslmaths "$ID"_rs_bold.nii.gz -mas "$ID"_rs_mask.nii.gz "$ID"_rs_bold_brain.nii.gz
#upscale voxel sizes
cp "$ID"_rs_bold_brain.nii.gz "$ID"_rs_bold_brain_x10.nii.gz
fslchpixdim "$ID"_rs_bold_brain_x10.nii.gz 3 10.5 3
rm *rs_t2*
fi
done
```

**g). Inspect the quality of reorientation and brain extraction (Quality control throughout step b-f)**

```
#!/bin/sh
module load fsl/6.0.3
mkdir -p brkraw_inspection/brain  brkraw_inspection/rs_brain
for file in PATH/TO/Bids_folder/sub-Cohort1*/ses*/anat/*brain.nii.gz; do
folder=${file%/*}
ID=${file##*/}
ID=${ID%".nii.gz"}  #get animal ID
echo $ID
slices "$file" "$folder"/"$ID"_T2w_coronal.nii.gz -o PATH/TO/brkraw_inspection/brain/"$ID"
done
for file in PATH/TO/Bids_folder/sub-Cohort1*/ses*/func/*brain_x10.nii.gz; do
ID=${file##*/}
ID=${ID%".nii.gz"}
echo $ID
slices "$file" -o PATH/TO/brkraw_inspection/rs_brain/"$ID"
done
```

**h). Denoise upscaled functional brain images using FSL/MELODIC and FIX**

Single-session ICA was first performed on the FSL/MELODIC GUI without a script, and then further de-noised using FIX with a script shown below.

```
#!/bin/sh
module load fsl/6.0.3
for folder in PATH/TO/Bids_folder/sub-Cohort*/ses*/func; do
        [ -d "$folder" ] && cd "$folder" &&
echo running FIX on "$folder"
/usr/local/fix/1.068/bin/fix sub-Cohort*.ica PATH/TO/FIX_training/FIX_training.RData 20
done
```

**i). Each de-noised functional brain image was registered to a Sprague-Dawley rat brain atlas**

```
#!/bin/sh
module load fsl/6.0.3
for folder in PATH/TO/Bids_folder/sub-Cohort*/ses*/func/*.ica; do
    [ -d "$folder" ] && cd "$folder" &&
ID=${folder#*func/}
ID=${ID%_x10*}
echo "doing coregistrations for "$ID""
flirt -in filtered_func_data_clean.nii.gz -ref PATH/TO/Atlas/atlasX10_downsampled8.nii.gz -out
"$ID"_clean_t2atlas.nii.gz -applyxfm -init reg/example_func2standard.mat -interp trilinear

mv filtered_func_data_clean.nii.gz "$ID"_clean.nii.gz
mv "$ID"_clean*.nii.gz ../
done
```

#### *B). Functional Component Dimensionality Optimization*

```
#!/bin/sh
module load fsl/6.0.3
#Based on the mICA toolbox, Sigma=fwhm/2.36
for folder in PATH/TO/Bids_folder/sub-Cohort*/ses*/func; do
        [ -d "$folder" ] && cd "$folder" &&
ID=${folder#*Bids_folder/}
ID=${ID%/func}
ID=${ID////_}

echo "spatial smoothing 3X altas on "$ID""
fslmaths "$ID"_rs_bold_brain_clean_t2atlas.nii.gz -mas
PATH/TO/Atlas/atlasX10_downsampled8_mask_GM.nii.gz -s 3.97 -mas
PATH/TO/Atlas/atlasX10_downsampled8_mask_GM.nii.gz
"$ID"_rs_bold_brain_clean_t2atlas_masked_s3.97.nii.gz

echo "spatial smoothing 4X altas on "$ID""
fslmaths "$ID"_rs_bold_brain_clean_t2atlas.nii.gz -mas
PATH/TO/Atlas/atlasX10_downsampled8_mask_GM.nii.gz -s 5.30 -mas
PATH/TO/Atlas/atlasX10_downsampled8_mask_GM.nii.gz
"$ID"_rs_bold_brain_clean_t2atlas_masked_s5.30.nii.gz

echo "spatial smoothing 5X altas on "$ID""
fslmaths "$ID"_rs_bold_brain_clean_t2atlas.nii.gz -mas
PATH/TO/Atlas/atlasX10_downsampled8_mask_GM.nii.gz -s 6.62 -mas
PATH/TO/Atlas/atlasX10_downsampled8_mask_GM.nii.gz
"$ID"_rs_bold_brain_clean_t2atlas_masked_s6.62.nii.gz
done

ls -1 PATH/TO/Bids_folder/sub-Cohort*/ses-Baseline/func/*masked_s6.62.nii.gz >
Analysis_ICA/All_Baseline_ICA_cleaned_s6.62.txt

cd PATH/TO/
python -m Analysis_ICA/mICA_Toolbox/py/splithalf Analysis_ICA/All_Baseline_ICA_cleaned_s6.62.txt
50 Analysis_ICA/FWHM_5Xatlas

cp -r Analysis_ICA/FWHM_5Xatlas Analysis_ICA/FWHM_4Xatlas
for file in Analysis_ICA/FWHM_4Xatlas/sample*/*.txt; do sed -i -e 's/s6.62/s5.30/g' "$file"; done

cp -r Analysis_ICA/FWHM_5Xatlas Analysis_ICA/FWHM_3Xatlas
for file in Analysis_ICA/FWHM_3Xatlas/sample*/*.txt; do sed -i -e 's/s6.62/s3.97/g' "$file"; done
```

**Continued**

```
#For parallel computation
for folder in PATH/TO/Analysis_ICA/FWHM_*Xatlas/sample*; do
    [ -d "$folder" ] && cd "$folder" &&
echo "$folder"
for dim in {10..300..10}; do
((i=i%12)); ((i++==0)) && wait
echo "$dim"
melodic -i group1_input.txt -o group1/dim"$dim" -v --nobet --bgthreshold=3 --tr=1.500000 --dim="$dim" --mmthresh=0.5 -m PATH/TO/Atlas/atlasX10_downsampled8_mask_GM.nii.gz
melodic -i group2_input.txt -o group2/dim"$dim"  -v --nobet --bgthreshold=3 --tr=1.500000 --dim="$dim" --mmthresh=0.5 -m PATH/TO/Atlas/atlasX10_downsampled8_mask_GM.nii.gz &
done
done

python mICA_Toolbox/py/ic_corr.py FWHM_5Xatlas 50 10,20,30,40,50,60,70,80,90,100,110,120,130,140,150,160,170,180,190,200,210,220,230,240,250,260,270,280,290,300 1

python mICA_Toolbox/py/ic_corr.py FWHM_4Xatlas 50 10,20,30,40,50,60,70,80,90,100,110,120,130,140,150,160,170,180,190,200,210,220,230,240,250,260,270,280,290,300 1

python mICA_Toolbox/py/ic_corr.py FWHM_3Xatlas 50 10,20,30,40,50,60,70,80,90,100,110,120,130,140,150,160,170,180,190,200,210,220,230,240,250,260,270,280,290,300 1
```

#### C). Reproducible Functional Component Identification

Several changes to gRAICAR codes (gRAICAR_step3.m) were done to match the spatial resolution (3.125 × 3.125 × 3.125 mm3) of our dataset

```
#!/bin/sh
module load fsl/6.0.3
module load matlab/r2019b
cd /PATH/TO/Analysis_ICA/RAICAR_reproducibility/FWHM_5Xatlas_dim220

#Make a list for all melodic_IC
ls sample*/group*/dim220/melodic_IC.nii.gz > FWHM_5Xatlas_dim220.list

#Create group mask
for i in {02..50}; do fslmaths sample_0001/group1/dim220/mask.nii.gz -mul
sample_00"$i"/group1/dim220/mask.nii.gz mask_FWHM_5Xatlas_dim220.nii.gz;done
for i in {01..50}; do fslmaths mask_FWHM_5Xatlas_dim220.nii.gz -mul
sample_00"$i"/group2/dim220/mask.nii.gz -bin mask_FWHM_5Xatlas_dim220.nii.gz;done

#Running gRAICAR using slurm
#!/bin/bash
#SBATCH --job-name=gRAICAR
#SBATCH --account=account_ID
#SBATCH --time=12:00:00
#SBATCH --nodes=1
#SBATCH --cpus-per-task=20
#SBATCH --gres=gpu:A40:1
#SBATCH --partition=gpu

module load fsl/6.0.3
module load matlab

cd PATH/TO/Analysis_ICA

matlab -nodisplay -nosplash -r "run('gRAICAR_5Xatlas_dim220.m');"
```

***Continued***

```
#Matlab script of 'gRAICAR_5Xatlas_dim220.m':
addpath (genpath('PATH/TO/Analysis_ICA/gRAICAR-master'));
addpath('PATH/TO/Analysis_ICA/FWHM_5Xatlas');
addpath('PATH/TO/Analysis_ICA');
settings.workdir = 'PATH/TO/Analysis_ICA';
settings.outdir = 'PATH/TO/Analysis_ICA/RAICAR_reproducibility/FWHM_5Xatlas_dim220';
settings.subjlist = 'PATH/TO/Analysis_ICA/FWHM_5Xatlas_dim220.list';
settings.maskpath = 'PATH/TO/Analysis_ICA/mask_FWHM_5Xatlas_dim220.nii.gz';
settings.taskname = 'FWHM_5Xatlas_dim220';
settings.ncores = 1;
settings.useRAICAR  = 0;
settings.icapath  =
'PATH/TO/Analysis_ICA/FWHM_5Xatlas/sample_0001/group1/dim220/melodic_IC.nii.gz';
settings.savemovie = 0;
settings.webreport = 1;
settings.displayThreshold = 1.5;
settings.comPerPage = 10;
[pass, settings] = gRAICAR_check_settings (settings);
[status, exeption] = gRAICAR_step1(settings);
[status, exeption] = gRAICAR_step2(settings);
[status, exeption] = gRAICAR_step3(settings);

##Back to terminal
module load convert3d
cd PATH/TO/Analysis_ICA/RAICAR_reproducibility
## for mixture modelling
echo "1" > grot.txt
fslmerge -t melodic_IC217.nii.gz ../FWHM_5Xatlas_dim220/compMaps/comp{001..217}.nii
melodic -i melodic_IC217.nii.gz --ICs=melodic_IC217.nii.gz --mix=grot.txt -o melodic217_Zstat --Oall --
report -v --mmthresh=0.5
# rename thresholded ICs using leading zeros to avoid wrong order
for i in {1..217}; do immv melodic217_Zstat/stats/thresh_zstat"${i}".nii.gz
melodic217_Zstat/stats/thresh_zstat`printf "%04g\n" $i`.nii.gz; done
fslmerge -t all_thresh_zstat217.nii.gz melodic217_Zstat/stats/thresh_zstat*.nii.gz
```

***Continued***

```
#nilearn for parcellation
source PATH/TO/Twainpythonenv3.8/bin/activate
ipython
import time
import matplotlib
import matplotlib.pyplot as plt
import numpy as np
from matplotlib import patches, ticker
import nilearn
from nilearn import datasets, plotting, image
from nilearn.image import get_data, index_img, mean_img,load_img
from nilearn.regions import Parcellations
data = nilearn.image.load_img("all_thresh_zstat217.nii.gz")
mask = nilearn.image.load_img("PATH/TO/Atlas/atlasX10_downsampled8_mask_GM.nii.gz")
ward = Parcellations(method="ward",n_parcels=217,mask = mask,
standardize=False,smoothing_fwhm=None,mask_strategy= "background",
memory="nilearn_cache",memory_level=1,verbose=1)
ward.fit(data)
ward_labels_img = ward.labels_img_
ward_labels_img.to_filename("ward_parcellation.nii.gz")

mkdir melodic217_Zstat/ward_mask
for i in {1..217}; do c3d ward_parcellation.nii.gz -thresh "$i" "$i" 1 0 -o
melodic217_Zstat/ward_mask/mask`printf "%04g\n" $i`.nii.gz;done

mkdir melodic217_Zstat/ward_node217
for n in {1..217}; do
((i=i%12)); ((i++==0)) && wait
echo $i
fslmaths melodic_IC217.nii.gz -Tmax -mas melodic217_Zstat/ward_mask/mask`printf "%04g\n"
$n`.nii.gz melodic217_Zstat/ward_node217/node`printf "%04g\n" $n`.nii.gz &
done

fslmerge -t ward_parcellation_IC217.nii.gz melodic217_Zstat/ward_node217/node*.nii.gz

slices_summary ward_parcellation_IC217 3
PATH/TO/Analysis_ICA/RAICAR_reproducibility/bgimage.nii.gz ward_parcellation_IC217.sum -1
```

#### D). Hierarchical Clustering

```
#!/bin/sh
module load fsl/6.0.3
module load matlab/r2019b
mkdir ts217
#extract time series of 217 nodes for all animals
for file in PATH/TO/Bids_folder/sub-*/ses-*/func/*_rs_bold_brain_clean_t2atlas_masked_s6.62.nii.gz;
do
((i=i%10)); ((i++==0)) && wait
ID=${file##*/}
ID=${ID%"_rs_bold_brain_clean_t2atlas_masked_s6.62.nii.gz"}
echo $ID
fsl_glm -i "$file" -d ward_parcellation_IC217 -o ts217/"$ID".txt --demean -m
PATH/TO/Atlas/atlasX10_downsampled8_mask_GM.nii.gz &
done

cd ts217
mkdir Baseline
for file in *Baseline.txt; do cp $file Baseline;done
mkdir All_timeseries
for file in *.txt; do mv $file All_timeseries;done
ls All_timeseries > All_timeseries_txt.csv

## Clustering in matlab
addpath PATH/TO/FSLNetwork_analysis/FSLNets;
addpath(sprintf('%s/usr/local/fsl/6.0.3/etc/matlab',getenv('fsl')));
addpath PATH/TO/Analysis_ICA/Auto-CVI-Tool/functions;
addpath PATH/TO/Analysis_ICA/Auto-CVI-Tool/functions/Hierarchichal;
addpath PATH/TO/Analysis_ICA/Auto-CVI-Tool/cvi;
addpath PATH/TO/Analysis_ICA/Auto-CVI-Tool/cvi/proximity;
addpath PATH/TO/Analysis_ICA/Auto-CVI-Tool/cvi/cvi_utils;

group_maps = 'ward_parcellation_IC217';
baseline_ts_dir = 'Baseline';
baseline_ts = nets_load(baseline_ts_dir, 1.5, 0);
baseline_ts.DD = [1:217];
baseline_Full = nets_netmats(baseline_ts,1,'corr');
[baseline_Znet_F,baseline_Mnet_F]=nets_groupmean(baseline_Full,1);
[Hier,Linkages] = nets_hierarchy(baseline_Znet_F,baseline_Znet_F,baseline_ts.DD,group_maps);

## Use Auto-CVI-Tool to determine the optimal number of networks
grot=prctile(abs(baseline_Znet_F(:)),99); netmatL=baseline_Znet_F/grot;
netmatH=baseline_Znet_F/grot;
usenet=netmatL;  usenet(usenet<0)=0;
N=size(netmatL,1);  grot=prctile(abs(usenet(:)),99); usenet=max(min(usenet/grot,1),-1)/2;
for J = 1:N, for I = 1:J-1,   y((I-1)*(N-I/2)+J-I) = 0.5 - usenet(I,J);  end; end;
y_reshape = squareform(y);
```

***Continued***

```
Kmax = 20;
clust = zeros(size(usenet,1),Kmax);
for k=1:Kmax
   clust(:,k) = cluster(Linkages, 'maxclust', k);
end

CVI = Select_CVI_Hierarchichal;
eva = evalcvi(clust,CVI,y_reshape);
Visualize_CVI(eva,CVI)

## Selectd CVI is:Index based on shapes, sizes, densities, and separation distances
writematrix(eva.FitnessValues, "optimal_CVI.csv")

Clusters = clust(:,13);
Seq = transpose(1:217);
Clusters_Seq = horzcat(Seq,Clusters);
Clusters_Seq = array2table(Clusters_Seq,'VariableNames',{'Node','Cluster'});

Hier_transpose = Hier';
Hier_transpose = array2table(Hier_transpose,'VariableNames',{'Node'});
Hier_Clusters = join(Hier_transpose,Clusters_Seq);
writetable(Hier_Clusters, "Node_Network13.xlsx")

## Extract full correlation based on re-ordered nodes
baseline_ts_reordered = baseline_ts;
baseline_ts_reordered.DD =
[186;195;177;10;1;135;167;31;11;69;26;34;41;46;130;204;4;21;125;56;122;58;100;113;183;8;53;80;111
;68;89;145;50;86;88;107;152;101;209;217;5;150;83;30;118;6;52;64;133;184;142;215;13;66;98;17;47;32
;129;123;18;162;70;212;191;193;214;39;45;165;180;27;148;166;28;178;176;168;196;37;205;211;42;14
4;57;202;158;9;23;112;126;90;138;156;207;12;131;172;110;154;132;49;149;55;54;140;170;93;146;147;
190;2;109;199;99;14;104;92;36;85;71;128;114;134;48;151;179;197;82;97;198;102;175;164;210;3;120;9
5;73;187;94;174;201;136;188;96;117;203;22;40;79;72;115;38;163;35;194;63;7;127;173;78;106;143;24;
139;59;108;169;15;105;16;20;33;77;43;61;171;19;65;81;206;116;124;137;213;161;182;76;160;91;157;1
89;153;155;185;25;103;192;119;121;159;29;208;62;200;44;75;74;87;51;67;84;181;60;216;141]';

baseline_ts_reordered = nets_tsclean(baseline_ts_reordered,0);
baseline_Full_reordered = nets_netmats(baseline_ts_reordered,1,'corr');
[baseline_reordered_Znet_F,baseline_reordered_Mnet_F]=nets_groupmean(baseline_Full_reordered,1
);
writematrix(baseline_reordered_Znet_F,"Baseline_reordered_Znet_Full.csv");

for n = 1:13
strcat("node=({",sprintf('%i,',table2array(Hier_Clusters(Hier_Clusters.Cluster == n,1))'),"})")
end
```

***Continued***

```
#create empty image for fsl/add
fslmaths ward_parcellation_IC217.nii.gz -Tstd -bin -thr 2 empty.nii.gz
for i in {1..13}; do cp empty.nii.gz Network`printf "%02g\n" $i`_add.nii.gz;done

#Network01 16 ICs
node1=({186,195,177,10,1,135,167,31,11,69,26,34,41,46,130,204})
echo "${node1[@]}"  ##list
echo "${#node1[@]}" ##count
node1=`printf "%04g\n" ${node1[@]}`
echo $node1 | sed 's/ /,/g'
for i in {186,195,177,10,1,135,167,31,11,69,26,34,41,46,130,204}; do echo "$i" && fslmaths
Network01_add.nii.gz -add melodic217_Zstat/ward_node217/node`printf "%04g\n" $i`
Network01_add.nii.gz;done
fslmerge -t Network01.nii.gz
melodic217_Zstat/ward_node217/node{0186,0195,0177,0010,0001,0135,0167,0031,0011,0069,0026,0
034,0041,0046,0130,0204}.nii.gz
#Network02 16 ICs
node2=({4,21,125,56,122,58,100,113,183,8,53,80,111,68,89,145})
node2=`printf "%04g\n" ${node2[@]}`
echo $node2 | sed 's/ /,/g'
for i in {4,21,125,56,122,58,100,113,183,8,53,80,111,68,89,145}; do echo "$i" && fslmaths
Network02_add.nii.gz -add melodic217_Zstat/ward_node217/node`printf "%04g\n" $i`
Network02_add.nii.gz;done
fslmerge -t Network02.nii.gz
melodic217_Zstat/ward_node217/node{0004,0021,0125,0056,0122,0058,0100,0113,0183,0008,0053,0
080,0111,0068,0089,0145}.nii.gz
#Network03 8 ICs
node3=({50,86,88,107,152,101,209,217})
node3=`printf "%04g\n" ${node3[@]}`
echo $node3 | sed 's/ /,/g'
for i in {50,86,88,107,152,101,209,217}; do echo "$i" && fslmaths Network03_add.nii.gz -add
melodic217_Zstat/ward_node217/node`printf "%04g\n" $i` Network03_add.nii.gz;done
fslmerge -t Network03.nii.gz
melodic217_Zstat/ward_node217/node{0050,0086,0088,0107,0152,0101,0209,0217}.nii.gz
#Network04 12 ICs
node4=({5,150,83,30,118,6,52,64,133,184,142,215})
node4=`printf "%04g\n" ${node4[@]}`
echo $node4 | sed 's/ /,/g'
for i in {5,150,83,30,118,6,52,64,133,184,142,215}; do echo "$i" && fslmaths Network04_add.nii.gz -
add melodic217_Zstat/ward_node217/node`printf "%04g\n" $i` Network04_add.nii.gz;done
fslmerge -t Network04.nii.gz
melodic217_Zstat/ward_node217/node{0005,0150,0083,0030,0118,0006,0052,0064,0133,0184,0142,0
215}.nii.gz
```

***Continued***

```
#Network05 8 ICs
node5=({13,66,98,17,47,32,129,123})
node5=`printf "%04g\n" ${node5[@]}`
echo $node5 | sed 's/ /,/g'
for i in {13,66,98,17,47,32,129,123}; do echo "$i" && fslmaths Network05_add.nii.gz -add
melodic217_Zstat/ward_node217/node`printf "%04g\n" $i` Network05_add.nii.gz;done
fslmerge -t Network05.nii.gz
melodic217_Zstat/ward_node217/node{0013,0066,0098,0017,0047,0032,0129,0123}.nii.gz
#Network06 11 ICs
node6=({18,162,70,212,191,193,214,39,45,165,180})
node6=`printf "%04g\n" ${node6[@]}`
echo $node6 | sed 's/ /,/g'
for i in {18,162,70,212,191,193,214,39,45,165,180}; do echo "$i" && fslmaths Network06_add.nii.gz -
add melodic217_Zstat/ward_node217/node`printf "%04g\n" $i` Network06_add.nii.gz;done
fslmerge -t Network06.nii.gz
melodic217_Zstat/ward_node217/node{0018,0162,0070,0212,0191,0193,0214,0039,0045,0165,0180}.n
ii.gz
#Network07 16 ICs
node7=({27,148,166,28,178,176,168,196,37,205,211,42,144,57,202,158})
node7=`printf "%04g\n" ${node7[@]}`
echo $node7 | sed 's/ /,/g'
for i in {27,148,166,28,178,176,168,196,37,205,211,42,144,57,202,158}; do echo "$i" && fslmaths
Network07_add.nii.gz -add melodic217_Zstat/ward_node217/node`printf "%04g\n" $i`
Network07_add.nii.gz;done
fslmerge -t Network07.nii.gz
melodic217_Zstat/ward_node217/node{0027,0148,0166,0028,0178,0176,0168,0196,0037,0205,0211,0
042,0144,0057,0202,0158}.nii.gz
#Network08 24 ICs
node8=({9,23,112,126,90,138,156,207,12,131,172,110,154,132,49,149,55,54,140,170,93,146,147,190})
node8=`printf "%04g\n" ${node8[@]}`
echo $node8 | sed 's/ /,/g'
for i in {9,23,112,126,90,138,156,207,12,131,172,110,154,132,49,149,55,54,140,170,93,146,147,190};
do echo "$i" && fslmaths Network08_add.nii.gz -add melodic217_Zstat/ward_node217/node`printf
"%04g\n" $i` Network08_add.nii.gz;done
fslmerge -t Network08.nii.gz
melodic217_Zstat/ward_node217/node{0009,0023,0112,0126,0090,0138,0156,0207,0012,0131,0172,0
110,0154,0132,0049,0149,0055,0054,0140,0170,0093,0146,0147,0190}.nii.gz
#Network09 24 ICs
node9=({2,109,199,99,14,104,92,36,85,71,128,114,134,48,151,179,197,82,97,198,102,175,164,210})
node9=`printf "%04g\n" ${node9[@]}`
echo $node9 | sed 's/ /,/g'
for i in {2,109,199,99,14,104,92,36,85,71,128,114,134,48,151,179,197,82,97,198,102,175,164,210}; do
echo "$i" && fslmaths Network09_add.nii.gz -add melodic217_Zstat/ward_node217/node`printf
"%04g\n" $i` Network09_add.nii.gz;done
fslmerge -t Network09.nii.gz
melodic217_Zstat/ward_node217/node{0002,0109,0199,0099,0014,0104,0092,0036,0085,0071,0128,0
114,0134,0048,0151,0179,0197,0082,0097,0198,0102,0175,0164,0210}.nii.gz
```

***Continued***

```
#Network10 23 ICs
node10=({3,120,95,73,187,94,174,201,136,188,96,117,203,22,40,79,72,115,38,163,35,194,63})
node10=`printf "%04g\n" ${node10[@]}`
echo $node10 | sed 's/ /,/g'
for i in {3,120,95,73,187,94,174,201,136,188,96,117,203,22,40,79,72,115,38,163,35,194,63}; do echo "$i" && fslmaths Network10_add.nii.gz -add melodic217_Zstat/ward_node217/node`printf "%04g\n" $i` Network10_add.nii.gz;done
fslmerge -t Network10.nii.gz melodic217_Zstat/ward_node217/node{0003,0120,0095,0073,0187,0094,0174,0201,0136,0188,0096,0117,0203,0022,0040,0079,0072,0115,0038,0163,0035,0194,0063}.nii.gz
#Network11 11 ICs
node11=({7,127,173,78,106,143,24,139,59,108,169})
node11=`printf "%04g\n" ${node11[@]}`
echo $node11 | sed 's/ /,/g'
for i in {7,127,173,78,106,143,24,139,59,108,169}; do echo "$i" && fslmaths Network11_add.nii.gz -add melodic217_Zstat/ward_node217/node`printf "%04g\n" $i` Network11_add.nii.gz;done
fslmerge -t Network11.nii.gz melodic217_Zstat/ward_node217/node{0007,0127,0173,0078,0106,0143,0024,0139,0059,0108,0169}.nii.gz
#Network12 27 ICs
node12=({15,105,16,20,33,77,43,61,171,19,65,81,206,116,124,137,213,161,182,76,160,91,157,189,153,155,185})
node12=`printf "%04g\n" ${node12[@]}`
echo $node12 | sed 's/ /,/g'
for i in {15,105,16,20,33,77,43,61,171,19,65,81,206,116,124,137,213,161,182,76,160,91,157,189,153,155,185}; do echo "$i" && fslmaths Network12_add.nii.gz -add melodic217_Zstat/ward_node217/node`printf "%04g\n" $i` Network12_add.nii.gz;done
fslmerge -t Network12.nii.gz melodic217_Zstat/ward_node217/node{0015,0105,0016,0020,0033,0077,0043,0061,0171,0019,0065,0081,0206,0116,0124,0137,0213,0161,0182,0076,0160,0091,0157,0189,0153,0155,0185}.nii.gz
#Network13 21 ICs
node13=({25,103,192,119,121,159,29,208,62,200,44,75,74,87,51,67,84,181,60,216,141})
node13=`printf "%04g\n" ${node13[@]}`
echo $node13 | sed 's/ /,/g'
for i in {25,103,192,119,121,159,29,208,62,200,44,75,74,87,51,67,84,181,60,216,141}; do echo "$i" && fslmaths Network13_add.nii.gz -add melodic217_Zstat/ward_node217/node`printf "%04g\n" $i` Network13_add.nii.gz;done
fslmerge -t Network13.nii.gz melodic217_Zstat/ward_node217/node{0025,0103,0192,0119,0121,0159,0029,0208,0062,0200,0044,0075,0074,0087,0051,0067,0084,0181,0060,0216,0141}.nii.gz

fslmerge -t Networks13.nii.gz Network{01..13}_add.nii.gz
```

***Continued***

```
mkdir Networks13
#extract time series of 13 networks for all animals
for file in PATH/TO/Bids_folder/sub-*/ses-*/func/*_rs_bold_brain_clean_t2atlas_masked_s6.62.nii.gz;
do
((i=i%10)); ((i++==0)) && wait
ID=${file##*/}
ID=${ID%"_rs_bold_brain_clean_t2atlas_masked_s6.62.nii.gz"}
echo $ID
fsl_glm -i "$file" -d Networks13.nii.gz -o Networks13/"$ID".txt --demean -m
PATH/TO/Atlas/atlasX10_downsampled8_mask_GM.nii.gz &
done

#extract volume for networks
for i in {01..13}; do echo "$i" && for mask in
PATH/TO/Atlas/WHS_SD_rat_lr_masks/unilateral/*_lrmask.nii.gz; do echo "$mask" &&
region=${mask%"_lrmask.nii.gz"} &&
region=${region#"PATH/TO/Atlas/WHS_SD_rat_lr_masks/unilateral/"} && stats=$(fslstats
Network"$i"_add.nii.gz -k "$mask" -M -V) && echo -e Network"$i" $region $stats >>
networks13_avgZ_volume.xls;done;done

cd melodic217_Zstat/ward_node217
mkdir Network{01..13}
for i in {186,195,177,10,1,135,167,31,11,69,26,34,41,46,130,204}; do echo "$i" && cp node`printf
"%04g\n" $i`.nii.gz Network01/;done

for i in {4,21,125,56,122,58,100,113,183,8,53,80,111,68,89,145}; do echo "$i" && cp node`printf
"%04g\n" $i`.nii.gz Network02/;done

for i in {50,86,88,107,152,101,209,217}; do echo "$i" && cp node`printf "%04g\n" $i`.nii.gz
Network03/;done

for i in {5,150,83,30,118,6,52,64,133,184,142,215}; do echo "$i" && cp node`printf "%04g\n" $i`.nii.gz
Network04/;done

for i in {13,66,98,17,47,32,129,123}; do echo "$i" && cp node`printf "%04g\n" $i`.nii.gz Network05/;done

for i in {18,162,70,212,191,193,214,39,45,165,180}; do echo "$i" && cp node`printf "%04g\n" $i`.nii.gz
Network06/;done

for i in {27,148,166,28,178,176,168,196,37,205,211,42,144,57,202,158}; do echo "$i" && cp node`printf
"%04g\n" $i`.nii.gz Network07/;done

for i in {9,23,112,126,90,138,156,207,12,131,172,110,154,132,49,149,55,54,140,170,93,146,147,190};
do echo "$i" && cp node`printf "%04g\n" $i`.nii.gz Network08/;done

for i in {2,109,199,99,14,104,92,36,85,71,128,114,134,48,151,179,197,82,97,198,102,175,164,210}; do
echo "$i" && cp node`printf "%04g\n" $i`.nii.gz Network09/;done
```

***Continued***

```
for i in {3,120,95,73,187,94,174,201,136,188,96,117,203,22,40,79,72,115,38,163,35,194,63}; do echo "$i" && cp node`printf "%04g\n" $i`.nii.gz Network10/;done

for i in {7,127,173,78,106,143,24,139,59,108,169}; do echo "$i" && cp node`printf "%04g\n" $i`.nii.gz Network11/;done

for i in {15,105,16,20,33,77,43,61,171,19,65,81,206,116,124,137,213,161,182,76,160,91,157,189,153,155,185}; do echo "$i" && cp node`printf "%04g\n" $i`.nii.gz Network12/;done

for i in {25,103,192,119,121,159,29,208,62,200,44,75,74,87,51,67,84,181,60,216,141}; do echo "$i" && cp node`printf "%04g\n" $i`.nii.gz Network13/;done

#extract node volume under each Network
for i in {01..13}; do echo Network"$i" && for node in Network"$i"/node*.nii.gz; do echo $node && nodename=${node%".nii.gz"} && for mask in PATH/TO/Atlas/WHS_SD_rat_lr_masks/unilateral/*_lrmask.nii.gz; do echo "$mask" && region=${mask%"_lrmask.nii.gz"} && region=${region#"PATH/TO/Atlas/WHS_SD_rat_lr_masks/unilateral/"} && stats=$(fslstats $node -k $mask -M -V) && echo -e Network"$i" $nodename $region $stats
>> ../../Nodes217_avgZ_volume.xls;done;done;done
```

#### *E). Functional Connectivity Computation*

```
#!/bin/bash

module load matlab
module load fsl/6.0.3
## For Connectivity between pairs of 13 Networks on matlab
matlab
addpath PATH/TO/FSLNetwork_analysis/FSLNets;
addpath(sprintf('%s/usr/local/fsl/6.0.3/etc/matlab',getenv('fsl')));
addpath PATH/TO/FSLNetwork_analysis/L1precision;
All_ts_dir = 'Networks13';
All_ts = nets_load(All_ts_dir, 1.5, 0);
All_ts.DD = [1:13];
All_Pridgep = nets_netmats(All_ts,0,'ridgep',0.1);
All_Pridgep_matrix=reshape(transpose(All_Pridgep),[13,13,192]);
no_sub = size(All_Pridgep_matrix,3);

for i = 1:no_sub
Pridgep_triled(:,:,i) = tril(All_Pridgep_matrix(:,:,i),-1);
G_Pridgep = graph(Pridgep_triled(:,:,i),'lower');
Pridgep_edgelist{i} = G_Pridgep.Edges;
All_edgestrength{i} = Pridgep_edgelist{i}.Weight';
End

All_Edgestrength = vertcat(All_edgestrength{:});
All_edgelist = Pridgep_edgelist{1,1}.EndNodes;
All_Edgelist = strcat("Network",num2str(All_edgelist(:,1)),"&",num2str(All_edgelist(:,2)));
All_Edgelist = All_Edgelist';
All_mat = vertcat(All_Edgelist,All_Edgestrength);

ID = importdata("All_timeseries_txt.csv");
match = ["sub-","ses-",".txt"];
ID = erase(ID,match);
ID = strrep(ID,"Cohort1-Control","Cohort1 Control");
ID = strrep(ID,"Cohort1-","Cohort1 CRS-");
ID = strrep(ID,"2-","2 ");
ID = strrep(ID,"3-","3 ");
ID = strrep(ID,"4-","4 ");
ID = strrep(ID,"-C"," C");
ID = strrep(ID,"_"," ");
ID = regexp(ID, ' ', 'split');
ID = vertcat(ID{:});
ID = table(ID);
ID = splitvars(ID,'ID','NewVariableNames',{'Cohort','Group','ID','Timepoint'});
ID.Cohort_ID = strcat(ID.Cohort, '_', ID.ID);
ID = movevars(ID,"Cohort_ID",'Before',"Cohort");
ID = movevars(ID,"Group",'After',"ID");
writetable(ID,"All_timeseries_metadata.csv");
```

***Continued***

```
ID = readtable("../All_timeseries_metadata.csv",'PreserveVariableNames', true);
ID(:, 2:3) = [];
ID1 = table2cell(ID);
ID1 = [ID.Properties.VariableNames;ID1];
All_mat = horzcat(ID1,All_mat);
writematrix(All_mat," 13Network_Connectivity.csv");
```

#### EPM Automation

Coordinates of body parts in each frame of each video were extracted using DeepLabCut GUI. These coord files were processed with DLCAnalyzer and and in-house R scripts in RStudio and scripts are shown below.

```
library(sf)
library(sp)
library(imputeTS)
library(ggplot2)
library(ggmap)
library(cowplot)
library(corrplot)
library(keras)
library(tensorflow)
library(xlsx)
library(data.table)
library(tidyverse)
library(dplyr)
library(plyr)
library(openxlsx)
setwd("PATH/TO/ EPM_analysis/")
source('PATH/TO/EPM_analysis/R/DLCAnalyzer_Functions_final.R')

input_folder <- "PATH/TO/EPM_all_csv/"
files <- list.files(input_folder)
files [1:6]
pipeline <- function(path)
{
  Tracking <- ReadDLCDataFromCSV(file = path, fps = 25)
  Tracking <- CalibrateTrackingData(Tracking, method = "distance",in.metric = 110, points = c ("tl", "br"))
  zoneinfo <- read.table("PATH/TO/EPM_analysis/EPM_zoneinfo.csv", sep = ";", header = T)
  Tracking <- AddZones(Tracking,zoneinfo)
  Tracking <- CleanTrackingData(Tracking, likelihoodcutoff = 0.95, existence.pol =
ScalePolygon(Tracking$zones$arena, 1.0))
  Tracking <- EPMAnalysis(Tracking, movement_cutoff = 4,integration_period = 5,points =
c("nose","headcentre", "earl", "earr", "neck", "bodycentre", "hipl","hipr","tailbase"), nosedips = FALSE)
  return(Tracking)
}

TrackingAll <- RunPipeline(files, input_folder, FUN = pipeline)

#Moving speed of the body center is the resluts we want to extract when using the DLCAnalyzer Report
Report <- MultiFileReport(TrackingAll)
write.xlsx(Report, file = "EPM_All_raw.xlsx", row.names = FALSE, col.names = TRUE)

save(TrackingAll, files, file = "TrackingAll.rda")
```

***Continued***

```
#In-house scripts (credit to Twain:)

load("TrackingAll.rda")
x <- NULL
y <- NULL
Coords <- NULL
Coords_list <- list()
Rat_polygon <- list()
Rat_sf_list <- list()
Left_closed <- list()
Right_closed<- list()
Top_open <- list()
Bottom_open<- list()
Left_closed_st <- list()
Right_closed_st <- list()
Top_open_st <- list()
Bottom_open_st <- list()

for (i in files){
  for (f in seq_along(TrackingAll[[i]]$frame)){
    x[[i]] <-
c(TrackingAll[[i]]$data[["neck"]]$x[f],TrackingAll[[i]]$data[["hipl"]]$x[f],TrackingAll[[i]]$data[["hipr"]]$x[f],TrackingAll[[i]]$data[["neck"]]$x[f])
    y[[i]] <-
c(TrackingAll[[i]]$data[["neck"]]$y[f],TrackingAll[[i]]$data[["hipl"]]$y[f],TrackingAll[[i]]$data[["hipr"]]$y[f],TrackingAll[[i]]$data[["neck"]]$y[f])
    Coords <- cbind(x[[i]],y[[i]])
    Coords_list[[i]][[f]] <- Coords
    Rat_sf_list[[i]][[f]] <- st_polygon(list(Coords))
    }}

for (i in files){
    Left_closed[[i]] <- as.matrix(TrackingAll[[i]]$zones$closed.left)
    Right_closed[[i]] <- as.matrix(TrackingAll[[i]]$zones$closed.right)
    Top_open[[i]] <- as.matrix(TrackingAll[[i]]$zones$open.top)
    Bottom_open[[i]] <- as.matrix(TrackingAll[[i]]$zones$open.bottom)
    Bottom_open[[i]] <- rbind(Bottom_open[[i]], Bottom_open[[i]][1:1,])
    Left_closed[[i]] <- rbind(Left_closed[[i]], Left_closed[[i]][1:1,])
    Right_closed[[i]] <- rbind(Right_closed[[i]], Right_closed[[i]][1:1,])
    Top_open[[i]] <- rbind(Top_open[[i]], Top_open[[i]][1:1,])
    Left_closed_st[[i]] <- st_polygon(list(Left_closed[[i]]))
    Right_closed_st[[i]] <- st_polygon(list(Right_closed[[i]]))
    Top_open_st[[i]] <- st_polygon(list(Top_open[[i]]))
    Bottom_open_st[[i]] <- st_polygon(list(Bottom_open[[i]]))
  }
```

***Continued***

```
## Compute frame counts for animal polygon inside each arm, will be used to estimate the total time in each arm
Left_closed_count <- list()
Right_closed_count <- list()
Top_open_count <- list()
Bottom_open_count <- list()

for (i in files){
   for (f in 1:length(Rat_sf_list[[i]])){
   Left_closed_count[[i]][[f]] <- st_within(Rat_sf_list[[i]][[f]], Left_closed_st[[i]], sparse = FALSE)
   Right_closed_count[[i]][[f]] <- st_within(Rat_sf_list[[i]][[f]], Right_closed_st[[i]], sparse = FALSE)
   Top_open_count[[i]][[f]] <- st_within(Rat_sf_list[[i]][[f]], Top_open_st[[i]], sparse = FALSE)
   Bottom_open_count[[i]][[f]] <- st_within(Rat_sf_list[[i]][[f]], Bottom_open_st[[i]], sparse = FALSE)
   }}
save(Left_closed_count, Right_closed_count, Top_open_count, Bottom_open_count, files, file = "zones_count.rda")

Left_closed_counts <- rbind.fill(lapply(Left_closed_count,function(y){as.data.frame(t(y),stringsAsFactors=FALSE)}))
Left_closed_counts <- cbind(files, Left_closed_counts)

Right_closed_counts <- rbind.fill(lapply(Right_closed_count,function(y){as.data.frame(t(y),stringsAsFactors=FALSE)}))
Right_closed_counts <- cbind(files, Right_closed_counts)

Top_open_counts <- rbind.fill(lapply(Top_open_count,function(y){as.data.frame(t(y),stringsAsFactors=FALSE)}))
Top_open_counts <- cbind(files, Top_open_counts)

Bottom_open_counts <- rbind.fill(lapply(Bottom_open_count,function(y){as.data.frame(t(y),stringsAsFactors=FALSE)}))
Bottom_open_counts <- cbind(files, Bottom_open_counts)

Left_closed_counts <- Left_closed_counts %>% add_column(Zone = "Left closed", .before = "V1")
Right_closed_counts <- Right_closed_counts %>% add_column(Zone = "Right closed", .before = "V1")
Top_open_counts <- Top_open_counts %>% add_column(Zone = "Top open", .before = "V1")
Bottom_open_counts <- Bottom_open_counts %>% add_column(Zone = "Bottom open", .before = "V1")

save(Left_closed_counts, Right_closed_counts, Top_open_counts, Bottom_open_counts, file = "zones_counts_files.rda")

Arena_counts <- as.data.frame(do.call(rbind,list(Left_closed_counts,Right_closed_counts, Top_open_counts, Bottom_open_counts)))

colnames(Arena_counts)[1] <- "Animal_Timepoint"
Arena_counts$Animal_Timepoint <- gsub("DLC.*$", "",Arena_counts$Animal_Timepoint)
```

***Continued***

```
openxlsx::write.xlsx(Arena_counts, file = "EPM_time_in_zones_logi.xlsx", rowNames = FALSE,
colNames = TRUE)

EPM <- openxlsx::read.xlsx("EPM_time_in_zones_logi.xlsx")
EPM[is.na(EPM)] <- FALSE

# Convert logical YES and NO to numeric of 1 and 0
sapply(3:ncol(EPM), function(i) {
  EPM[, i] <<- as.numeric(EPM[, i])
})
openxlsx::write.xlsx(EPM, file = "EPM_time_in_zones.xlsx", rowNames = FALSE, colNames = TRUE)

## Compute frame counts for animal polygon outside each arm will be used to calculate the transition
across open and closed arms

Left_closed_rest <- list()
Right_closed_rest <- list()
Top_open_rest <- list()
Bottom_open_rest <- list()
Left_closed_rest_st <- list()
Right_closed_rest_st <- list()
Top_open_rest_st <- list()
Bottom_open_rest_st <- list()

for (i in files){
  arena_temp <- as.matrix(TrackingAll[[i]]$zones$arena)
  Left_closed_rest[[i]] <- arena_temp[!(row.names(arena_temp) %in% c("lb","lt")),]
  Left_closed_rest[[i]] <- rbind(Left_closed_rest[[i]], Left_closed_rest[[i]][1:1,])
  Right_closed_rest[[i]] <- arena_temp[!(row.names(arena_temp) %in% c("rb","rt")),]
  Right_closed_rest[[i]] <- rbind(Right_closed_rest[[i]], Right_closed_rest[[i]][1:1,])
  Top_open_rest[[i]] <- arena_temp[!(row.names(arena_temp) %in% c("tl","tr")),]
  Top_open_rest[[i]] <- rbind(Top_open_rest[[i]], Top_open_rest[[i]][1:1,])
  Bottom_open_rest[[i]] <- arena_temp[!(row.names(arena_temp) %in% c("bl","br")),]
  Bottom_open_rest[[i]] <- rbind(Bottom_open_rest[[i]], Bottom_open_rest[[i]][1:1,])
  Left_closed_rest_st[[i]] <- st_polygon(list(Left_closed_rest[[i]]))
  Right_closed_rest_st[[i]] <- st_polygon(list(Right_closed_rest[[i]]))
  Top_open_rest_st[[i]] <- st_polygon(list(Top_open_rest[[i]]))
  Bottom_open_rest_st[[i]] <- st_polygon(list(Bottom_open_rest[[i]]))
}

Left_closed_rest_count <- list()
Right_closed_rest_count <- list()
Top_open_rest_count <- list()
Bottom_open_rest_count <- list()
```

***Continued***

```
for (i in files){
  for (f in 1:length(Rat_sf_list[[i]])){
    Left_closed_rest_count[[i]][[f]] <- st_within(Rat_sf_list[[i]][[f]], Left_closed_rest_st[[i]], sparse =
FALSE)
    Right_closed_rest_count[[i]][[f]] <- st_within(Rat_sf_list[[i]][[f]], Right_closed_rest_st[[i]], sparse =
FALSE)
    Top_open_rest_count[[i]][[f]] <- st_within(Rat_sf_list[[i]][[f]], Top_open_rest_st[[i]], sparse = FALSE)
    Bottom_open_rest_count[[i]][[f]] <- st_within(Rat_sf_list[[i]][[f]], Bottom_open_rest_st[[i]], sparse =
FALSE)
  }}

save(Left_closed_rest_count, Right_closed_rest_count, Top_open_rest_count,
Bottom_open_rest_count, files, file = "zones_rest_count.rda")

Left_closed_rest_counts <-
rbind.fill(sapply(Left_closed_rest_count,function(y){as.data.frame(t(y),stringsAsFactors=FALSE)}))
Left_closed_rest_counts <- as.data.frame(cbind(files, Left_closed_rest_counts))

Right_closed_rest_counts <-
rbind.fill(lapply(Right_closed_rest_count,function(y){as.data.frame(t(y),stringsAsFactors=FALSE)}))
Right_closed_rest_counts <- as.data.frame(cbind(files, Right_closed_rest_counts))

Top_open_rest_counts <-
rbind.fill(lapply(Top_open_rest_count,function(y){as.data.frame(t(y),stringsAsFactors=FALSE)}))
Top_open_rest_counts <- as.data.frame(cbind(files, Top_open_rest_counts))

Bottom_open_rest_counts <-
rbind.fill(lapply(Bottom_open_rest_count,function(y){as.data.frame(t(y),stringsAsFactors=FALSE)}))
Bottom_open_rest_counts <- as.data.frame(cbind(files, Bottom_open_rest_counts))

  Left_closed_rest_counts <- Left_closed_rest_counts %>% add_column(Zone = "Left closed
rest", .before = "V1")
  Right_closed_rest_counts <- Right_closed_rest_counts %>% add_column(Zone = "Right closed
rest", .before = "V1")
  Top_open_rest_counts <- Top_open_rest_counts %>% add_column(Zone = "Top open rest", .before
= "V1")
  Bottom_open_rest_counts <- Bottom_open_rest_counts %>% add_column(Zone = "Bottom open
rest", .before = "V1")

  save(Left_closed_rest_counts, Right_closed_rest_counts, Top_open_rest_counts,
Bottom_open_rest_counts, file = "zones_rest_counts_files.rda")

Arena_rest_counts <-
as.data.frame(do.call(rbind,list(Left_closed_rest_counts,Right_closed_rest_counts,
Top_open_rest_counts, Bottom_open_rest_counts)))

colnames(Arena_rest_counts)[1] <- "Animal_Timepoint"
```

***Continued***

```
Arena_rest_counts$Animal_Timepoint <- gsub("DLC.*$", "",Arena_rest_counts$Animal_Timepoint)

  openxlsx::write.xlsx(Arena_rest_counts, file = "EPM_time_in_zones_rest_logi.xlsx", rowNames =
FALSE, colNames = TRUE)

  EPM_rest <- openxlsx::read.xlsx("EPM_time_in_zones_rest_logi.xlsx")

  EPM_rest[is.na(EPM_rest)] <- FALSE

 sapply(3:ncol(EPM_rest), function(i) {
   EPM_rest[, i] <<- as.numeric(EPM_rest[, i])
  })
 EPM_rest_neg <- EPM_rest
 EPM_rest_neg[sapply(EPM_rest_neg, is.numeric)] <- EPM_rest_neg[sapply(EPM_rest_neg,
is.numeric)] * -1

 openxlsx::write.xlsx(EPM_rest, file = "EPM_time_in_zones_rest.xlsx", rowNames = FALSE, colNames
= TRUE)
 openxlsx::write.xlsx(EPM_rest_neg, file = "EPM_time_in_zones_rest_negative.xlsx", rowNames =
FALSE, colNames = TRUE)

## Extract moving speed from results from Report generated in page 19, and calculate time spent in
each arm and transitions.

EPM_time_in_zones <- openxlsx::read.xlsx("EPM_time_in_zones.xlsx")
EPM_time_in_zones_rest <- openxlsx::read.xlsx("EPM_time_in_zones_rest_negative.xlsx")
EPM_raw <- openxlsx::read.xlsx("EPM_All_raw.xlsx")
EPM_raw <- EPM_raw %>% select(-contains(c('closed','open')))
EPM_raw <- EPM_raw %>% select(c(file, starts_with("bodycentre")))
EPM_raw <- EPM_raw[,-10:-18]
EPM_raw <- EPM_raw %>% select(c(file, contains(c("total.time","speed.moving"))))
EPM_raw <- EPM_raw %>% rename(Animal_Timepoint = 1, Total_time = 2,Moving_speed = 3)
EPM_raw$Animal_Timepoint <- gsub("DLC.*$", "",EPM_raw$Animal_Timepoint)

EPM_time_in_zones$Time_spent <- rowSums(EPM_time_in_zones == 1)
EPM_time_in_zones <- EPM_time_in_zones %>% relocate(Time_spent, .before = "V1")
EPM_time_frame_counts <- EPM_time_in_zones[1:3]
EPM_time_in_zones <- EPM_time_in_zones[,-3]
EPM_time_frame_counts <- spread(EPM_time_frame_counts, Zone, Time_spent)
EPM_time_frame_counts$Closed_frames <- (EPM_time_frame_counts$`Left closed` +
EPM_time_frame_counts$`Right closed`)
EPM_time_frame_counts$Open_frames <- (EPM_time_frame_counts$`Bottom open` +
EPM_time_frame_counts$`Top open`)
colnames(EPM_time_frame_counts)[c(2:5)] <- paste(colnames(EPM_time_frame_counts)[c(2:5)],
'frames', sep = " ")
```

***Continued***

```
#Calculate transitions across open and closed arms
EPM_time_in_zones <- EPM_time_in_zones %>% add_column(Part = "Main", .before = "V1")
EPM_time_in_zones_rest <- EPM_time_in_zones_rest  %>% add_column(Part = "Rest", .before =
"V1")
EPM_time_in_zones_rest <- data.frame(lapply(EPM_time_in_zones_rest, function(x){gsub(" rest", "",
x)}))
EPM_time_in_zones_rest <- EPM_time_in_zones_rest %>%
  mutate_at(c(4:7523), as.numeric) %>% mutate_at(c(1:3), as.factor)

EPM_logi <- rbind(EPM_time_in_zones, EPM_time_in_zones_rest)

EPM_logi_sum <- EPM_logi %>% as.data.frame() %>%
  group_by(Animal_Timepoint,Zone) %>%
    summarise(across(where(is.numeric), sum))

counts <- lapply(3:(ncol(EPM_logi_sum)-1), function(i){EPM_logi_sum[,i] + EPM_logi_sum[,i+1]})
counts_new <- do.call(cbind,counts)
counts_new <- cbind(EPM_logi_sum[1:2], counts_new)

transition_count <- list()
for(i in 1:nrow(counts_new)) {       # for-loop over rows
  data <- as.matrix(counts_new[i, ])
  data <- transpose(as.data.frame(as.numeric(data[1,data == 1 | data == -1])))
  if (length(data) <=1) {transition_count[[i]] <- 0
  } else {
  data_sum <- lapply(1:(length(data)-1), function(x){data[x] + data[x+1]})
  data_sum <- data.frame(do.call(cbind,data_sum))
  col_odd <- seq_len(ncol(data_sum)) %% 2
  data_sum <- data_sum[,col_odd == 1, drop = FALSE]
  transition_count[[i]] <- rowSums(data_sum == 0)
}}

Transitions <- do.call(rbind,transition_count)
Transitions <- cbind(EPM_logi_sum[1:2], Transitions)

colnames(Transitions)[3] <- "Transitions"
Transitions <- spread(Transitions, Zone, Transitions)

Transitions$Closed_transitions <- (Transitions$`Left closed` + Transitions$`Right closed`)
Transitions$Open_transitions <- (Transitions$`Bottom open` + Transitions$`Top open`)

colnames(Transitions)[c(2:5)] <- paste(colnames(Transitions)[c(2:5)], 'transitions', sep = " ")
```

***Continued***

```
##Combine moving speed, transitions and frames counts of time spent in each arm into one dataframe
df_list <- list(EPM_raw, Transitions,EPM_time_frame_counts)

EPM <- df_list  %>% reduce(full_join, "Animal_Timepoint")
EPM $Total_frames <- (EPM$Total_time * 25)
EPM$Percentage_closed_time <- (EPM$Closed_frames / EPM$Total_frames)
EPM$Percentage_open_time <- (EPM$Open_frames / EPM$Total_frames)

EPM <- EPM %>% mutate_if(is.numeric, round, 4)
EPM <- EPM %>% relocate(Percentage_closed_time, .after = "Total_time") %>%
  relocate(Percentage_open_time, .after = "Percentage_closed_time")%>%
  relocate(Closed_transitions, .before = "Total_time")%>%
  relocate(Open_transitions, .before = "Total_time")%>%
  relocate(Total_time, .after = "Total_frames")

EPM <- EPM %>% separate(Animal_Timepoint, c('Cohort', 'ID', 'Timepoint'), sep = "_", extra = "merge",
fill = "left") %>% unite("Cohort_ID", Cohort:ID, sep = "_", remove = FALSE) %>%
relocate(Cohort_ID, .before = "Cohort") %>% unite("Cohort_ID_Timepoint", Cohort:Timepoint, sep =
"_", remove = FALSE) %>% relocate(Cohort_ID_Timepoint, .before = "Cohort_ID") %>%
mutate(Timepoint = str_replace(Timepoint, "Depression","Post-CRS"))

xlsx::write.xlsx(as.data.frame(EPM), file = "EPM.xlsx", sheetName = "EPM", col.names = TRUE,
row.names = FALSE, append = TRUE)

## Note only moving speed, percentage of time spent in closed arms and transitions across closed
arms in the EPM test were later used in brain-behaviour modelling
```

#### Brain-Behaviour Modelling

```
module load R
R --max-ppsize 500000
## This R script is written by Twain Dai.
rm(list = ls())
.libPaths("/PATH/TO/R_lib")
library(dabestr)
library(tidyverse)
library(dplyr)
library(tidyr)
library(openxlsx)
library(str2str)
library(R.utils)
library(data.table)
library(parallel)
library(glmnet)
library(foreach)
require(vip)
library(caret)
library(ggalluvial)
library(smotefamily)
library(nestedcv)
library(Boruta)
library(pROC)
library(gapminder)

setwd("PATH/TO/Folder")
## Modelling with 13 Networks
#Load connectivity data from pairs of 13 networks and organise
Conn_CRS_BvD <- openxlsx::read.xlsx("13Network_Connectivity.xlsx")
COLNAMES <- data.frame(colnames(Conn_CRS_BvD))
colnames(COLNAMES)[1] <- "psudo"
COLNAMES <- COLNAMES %>% mutate(psudo = str_replace(psudo, "Network","Network.")) %>%
separate(psudo, c("Network", "V1", "V2")) %>%
  unite("Edge", Network:V2, sep = "_") %>%
  mutate(Edge = str_replace(Edge, "_NA","")) %>%
  mutate(Edge = str_replace(Edge, "_NA","")) %>% transpose()
COLNAMES <- as.data.frame(do.call(cbind,COLNAMES))
rownames(COLNAMES) <- NULL
colnames(Conn_CRS_BvD) <- unlist(COLNAMES)
Conn_CRS_BvD <- Conn_CRS_BvD %>%
  mutate_at(c(3:ncol(.)), as.numeric)

Conn_CRS_Baseline <- Conn_CRS_BvD %>%
   filter(Timepoint %in% "Baseline") %>% select(-c(Timepoint)) %>%
   rename_with(.fn = function(.x){paste0(.x,"_Baseline")},.cols=Network_1_2:Network_12_13)
```

***Continued***

```
Conn_CRS_diff <- Conn_CRS_BvD %>% arrange(Cohort_ID,Timepoint) %>%
  group_by(Cohort_ID) %>%
  summarise(mutate(across(Network_1_2:Network_12_13, function(x) {x[2]-x[1]}))) %>%
  rename_with(.fn = function(.x){paste0(.x,"_diff")},.cols=Network_1_2:Network_12_13)

#Load EPM data and organise
EPM_CRS_BvD <- openxlsx::read.xlsx("EPM.xlsx") %>%
  arrange(Cohort_ID,Timepoint)  %>%
  mutate(Anxiety_score = case_when(
    Percentage_closed_time >= 0.90 ~ 2,
    Percentage_closed_time >= 0.65 & Percentage_closed_time < 0.9 ~ 1,
    Percentage_closed_time < 0.65 ~ 0))

EPM_CRS_diff <- EPM_CRS_BvD %>% arrange(Cohort_ID,Timepoint) %>%
  group_by(Cohort_ID) %>% summarise(mutate(across(Percentage_closed_time:Anxiety_score,
function(x) {x[2]-x[1]}))) %>%  rename_with( .fn = function(.x){paste0(.x,"_Diff")},.cols=c(2:3))

df_list <- list(EPM_CRS_diff, Conn_CRS_diff[,1])
EPM_CRS_diff <- df_list %>% reduce(inner_join, c("Cohort_ID"))

EPM_CRS_BvD <- EPM_CRS_BvD %>%
  mutate(Anxiety = case_when(
    Percentage_closed_time >= 0.90 ~ "High Anxiety",
    Percentage_closed_time >= 0.65 & Percentage_closed_time < 0.9 ~ "Medium Anxiety",
    Percentage_closed_time < 0.65 ~ "Low Anxiety")
  ) %>%   filter(Cohort_ID %in% EPM_CRS_diff$Cohort_ID)

EPM_CRS_BvD_wide <- EPM_CRS_BvD[-2] %>%
  gather(key, value, -Cohort_ID, -Timepoint) %>%
  unite(col, key, Timepoint) %>%
  spread(col, value)  %>%
  mutate(Changes = case_when(
    Anxiety_PostCRS %in% "High Anxiety" & Anxiety_Baseline %in% "High Anxiety" ~ "High2High",
    Anxiety_PostCRS %in% "Medium Anxiety" & Anxiety_Baseline %in% "High Anxiety" ~
"High2Medium",
    Anxiety_PostCRS %in% "High Anxiety" & Anxiety_Baseline %in% "Medium Anxiety" ~
"Medium2High",
    Anxiety_PostCRS %in% "Medium Anxiety" & Anxiety_Baseline %in% "Medium Anxiety" ~
"Medium2Medium",
    Anxiety_PostCRS %in% "Low Anxiety" & Anxiety_Baseline %in% "Medium Anxiety" ~
"Medium2Low",
    Anxiety_PostCRS %in% "Medium Anxiety" & Anxiety_Baseline %in% "Low Anxiety" ~
"Low2Medium",
    Anxiety_PostCRS %in% "Low Anxiety" & Anxiety_Baseline %in% "Low Anxiety" ~ "Low2Low"
  )) %>% cbind(EPM_CRS_diff[-1])
```

***Continued***

```
#Classify animals into resilient and vulnerable to developing anxiety related behavior in EPM
Anxiety_changes <- as.data.frame(table(EPM_CRS_BvD_wide$Changes)) %>%
  mutate(Var1 = factor(Var1, levels = c("High2High",
"High2Medium","Medium2High","Medium2Medium","Medium2Low","Low2Medium", "Low2Low"))) %>%
arrange(Var1)

Anxiety_vulnerable <- EPM_CRS_BvD_wide %>%
  filter(Percentage_closed_time_Diff > 0) %>%
  filter(! (Anxiety_score_PostCRS == 0 & Anxiety_score_Baseline == 0)) %>%
  filter(! Anxiety_score_Baseline == 2) %>%
  add_column(Type = "Vulnerable") %>%
  relocate(Type, .after = "Cohort_ID")

Anxiety_resilience <- EPM_CRS_BvD_wide %>%
  filter(!Cohort_ID %in% Anxiety_vulnerable$Cohort_ID) %>%
  filter(!Anxiety_score_Baseline == 2)  %>%
  add_column(Type = "Resilience") %>%
  relocate(Type, .after = "Cohort_ID")

Anxiety <- rbind(Anxiety_vulnerable,Anxiety_resilience) %>%
  select(-Anxiety_Baseline,-contains("Anxiety_score"),-Changes,
        -Percentage_closed_time_PostCRS, -Anxiety_PostCRS)

#Load FST data and organise
FST_CRS_BvD <- openxlsx::read.xlsx("FST.xlsx") %>%
  mutate(Activity_Score = Swimming + Climbing - Immobility)

FST_CRS_diff <- FST_CRS_BvD[,c(1,3,8)] %>% arrange(Cohort_ID,Timepoint) %>%
  group_by(Cohort_ID) %>% summarise(mutate(across(Activity_Score, function(x) {x[2]-x[1]}))) %>%
  rename_with( .fn = function(.x){paste0(.x,"_Diff")},.cols=c(2))

df_list <- list(FST_CRS_diff, Conn_CRS_diff[,1])
FST_CRS_diff <- df_list %>% reduce(inner_join, c("Cohort_ID"))

FST_CRS_BvD <- FST_CRS_BvD  %>%  filter(Cohort_ID %in% FST_CRS_diff$Cohort_ID) %>%
  mutate(Activity = case_when(Activity_Score > 0 ~ "Active",
                              Activity_Score < 0 ~ "Passive",
                              TRUE ~ "Neutral"))
FST_CRS_BvD_wide <- FST_CRS_BvD[,c(1,3,8,9)] %>%
  filter(Cohort_ID %in% FST_CRS_diff$Cohort_ID) %>%
  gather(key, value, -Cohort_ID, -Timepoint) %>%
  unite(col, key, Timepoint) %>%
  spread(col, value) %>%
  mutate_at(c(1:3), factor) %>%
  mutate_at(c(4:5), as.numeric)
```

***Continued***

```
#Classify animals into resilient and vulnerable to developing passive coping behavior in FST
FST_CRS_BvD_wide <- FST_CRS_BvD_wide %>%
  cbind(FST_CRS_diff[,-1]) %>%
  mutate(Changes = case_when(
    Activity_PostCRS %in% "Active" & Activity_Baseline %in% "Active" ~ "A2A",
    Activity_PostCRS %in% "Neutral" & Activity_Baseline %in% "Active" ~ "A2N",
    Activity_PostCRS %in% "Passive" & Activity_Baseline %in% "Active" ~ "A2P",
    Activity_PostCRS %in% "Active" & Activity_Baseline %in% "Passive" ~ "P2A",
    Activity_PostCRS %in% "Passive" & Activity_Baseline %in% "Passive" ~ "P2P"))

Activity_changes <- as.data.frame(table(FST_CRS_BvD_wide$Changes)) %>%
  mutate(Var1 = factor(Var1, c("A2A","A2N","A2P","P2A","P2P"))) %>% arrange(Var1)

FST_vulnerable <- FST_CRS_BvD_wide %>%
  filter(Activity_Score_Diff < 0) %>%
  filter(! Activity_Baseline == "Passive")  %>%
  add_column(Type = "Vulnerable") %>%
  relocate(Type, .after = "Cohort_ID")

FST_resilience <- FST_CRS_BvD_wide %>%
  filter(!Cohort_ID %in% FST_vulnerable$Cohort_ID) %>%
  filter(! Activity_Baseline == "Passive")  %>%
  add_column(Type = "Resilience") %>%
  relocate(Type, .after = "Cohort_ID")

Activity <- rbind(FST_vulnerable,FST_resilience) %>%
  select(-Activity_Baseline,-Activity_PostCRS,-Activity_Score_PostCRS,-Changes)

#Combine connectivity and EPM data
df_list <- list(Anxiety,Conn_CRS_diff,Conn_CRS_Baseline)
Conn_Anxiety_Basediff <- df_list %>%
  reduce(inner_join, c("Cohort_ID")) %>%
  mutate(Type = as.factor(Type)) %>%
  mutate_at(c(3:4), as.numeric)

#Combine connectivity and FST data
df_list <- list(Activity,Conn_CRS_diff,Conn_CRS_Baseline)
Conn_Activity_Basediff <- df_list %>%
  reduce(inner_join, c("Cohort_ID")) %>%
  mutate(Type = as.factor(Type))  %>%
  mutate_at(c(3:4), as.integer)
```

```
#Perform nested cross validation 50 times for neurobehavior modeling using EPM data; the code for
modeling with FST data is same as follows.
Anxiety_repnCV <- foreach(i = 1:50,.packages=c('tidyverse','nestedcv','caret'),
                          .errorhandling = "remove") %dopar% {
                            nestcv.train(
                              y = Conn_Anxiety_Basediff$Type,
                              x = Conn_Anxiety_Basediff[,-1:-4],
                              method = "svmRadial",
                              balance = smote,
                              balance_options = list(k = 5),
                              filterFUN = boruta_filter,
                              filter_options = list(type = "names", select = "Confirmed",mcAdj = TRUE),
                              outer_method = "cv",
                              n_outer_folds = 5,
                              n_inner_folds = 4,
                              cv.cores = 2,
                              trControl = trainControl(classProbs = TRUE,
                                                       summaryFunction = twoClassSummary),
                              tuneLength = 10,
                              savePredictions = "final",
                              outer_train_predict = TRUE,
                              finalCV = TRUE
                            )
                          }
## Extract results from 50 repetitions of nested cross validation
Anxiety_Summary <- foreach(i = 1:50,.packages=c('tidyverse')) %do% {
  summary(Anxiety_repnCV[[1]])}

Anxiety_ModelSum <- foreach(i = 1:50,.packages=c('tidyverse'), .combine = rbind) %do% {
    data.frame(t((Anxiety_Summary[[i]]$result$metrics)))   %>%
    add_column(Predictors = length(Anxiety_repnCV[[i]]$final_vars)) %>%
    data.frame(t(confusionMatrix(Anxiety_Summary[[i]][["result"]][["table"]])$byClass)) %>%
    select(AUC,Balanced.Accuracy,Sensitivity,Specificity,F1,Predictors) %>%
    mutate_at("Predictors", as.integer)
}

Anxiety_Performance <-Anxiety_ModelSum %>% pivot_longer(cols = everything()) %>%
  group_by(name) %>%
  summarise(mean=mean(value), sd=sd(value), median=median(value),
          max=max(value), min=min(value))
```

```
##Extract variable stability measures
Anxiety_Varstability <- foreach(i = 1:50,.packages=c('tidyverse'), .combine = rbind) %do% {
  data.frame(var_stability(Anxiety_repnCV[[i]])) %>%
    add_column(Variables = row.names(.)) %>%
    filter(final == "yes") %>%
    add_column(Model = i)
}

## Extract variable importance and combine with stability measures
Anxiety_VarsImp <- Anxiety_Varstability %>%
    group_by(Variables) %>%
    summarise(Importance_mean=mean(mean), Importance_sd=sd(mean))

Anxiety_Vars <- table(Anxiety_Varstability$Variables) %>% as.data.frame() %>%
  `colnames<-`(c("Variables", "Frequency")) %>% arrange(desc(Frequency)) %>%
   inner_join(Anxiety_Varstability2[5:8], "Variables") %>% distinct()

df_list <- list(Anxiety_Vars,Anxiety_VarsImp)
Anxiety_Vars <-  df_list %>%
  reduce(inner_join, c("Variables")) %>%
   mutate(Importance_mean = Importance_mean * sign) %>%
   rename(Direction = direction) %>%
   mutate(Direction = case_when(
     Direction == "Up in Resilience" ~ "Up in Resilient",
     TRUE ~ "Up in Vulnerable"))
```

### Additional bash script for creating high and low resolution of atlas mask for each brain substructures with the Waxholm Space atlas (RRID:SCR_017124)

```
#!/bin/sh

module load convert3d
module load fsl/6.0.3
## This workflow of making atlas mask for brain substructures is credit to Twain Dai
.
## The software (fsl and convert3d) used in the workflow are credit to the original production teams.

## create pairs of substructure ID and their abbreviation
PAIRS=("410 AId" "424 AIp" "409 AIv" "501 Amu" "223 Ang" "213 AD" "254 AM" "214 AVdm" "215
AVvl" "82 BFRu" "93 BNST" "47 BSu" "197 CPu" "248 CL" "247 CM" "5 Cbu" "411 Cg1" "10 Cg2" "412
CLA" "98 CA1" "97 CA2" "95 CA3" "55 SuD" "96 DG" "128 DCND" "127 DCNFG" "126 DCNM" "205
DLG" "404 DLO" "414 DI" "500 Endo" "32 EP" "282 Eth" "99 FC" "280 FoF" "407 Fr3" "77 FrA" "198
GPel" "195 GPem" "64 GlA" "65 Gl" "416 GI" "48 HThu" "143 CNIC" "142 DCIC" "145 ECIC" "74 IO"
"413 IL" "255 IAM" "272 IGL" "260 IMD" "71 IP" "115 LEC" "206 LHb" "139 DLL" "138 ILL" "137 VLL"
"401 LO" "283 LPl" "286 LPmc" "285 LPmr" "134 LSO" "228 LDdm" "229 LDvl" "114 MEC" "295 MGd"
"150 MGmz" "297 MGm" "299 MGsg" "298 MGv" "207 MHb" "403 MO" "133 MSO" "233 MDc" "232
MDl" "240 MDm" "4 Cbm" "184 NAcc" "192 NAcsh" "502 NLOT" "81 SMn" "131 NTB" "163 Sag" "66
OBu" "246 PCN" "267 PF" "110 PaS" "211 PT" "242 PV" "432 lPPC" "433 mPPC" "436 PtP" "51 PAG"
"201 PP" "112 PER35" "113 PER36" "56 PVG" "43 PG" "181 PIR1" "182 PIR2" "183 PIR3" "58 Pn" "208
PIL" "210 Pot" "230 Po" "108 POR" "204 PrG" "405 PrL" "109 PrS" "94 PRT" "151 Au1" "408 M1" "425
S1bf" "418 S1dz" "420 S1f" "417 S1fl" "423 S1hl" "429 S1tr" "442 V1" "164 RTa" "200 RTu" "268 RRe"
"427 RSD" "430 RSG" "219 Re" "216 Rh" "152 Au2d" "153 Au2v" "406 M2" "422 S2" "443 V2L" "448
V2M" "40 Sep" "75 Sp5n" "281 SubG" "100 SUB" "222 SMT" "278 SPF" "188 SNc" "189 SNl" "187 SNr"
"3 STh" "50 SuG" "132 SPN" "135 SPR" "444 TeA" "293 VA" "158 AVCN" "160 Cap" "123 GCL" "159
PVCN" "402 VO" "193 VP" "136 VPO" "266 VPpc" "294 VPL" "227 VPM" "199 VSRu" "196 VTA" "400
VLO" "231 VL" "221 VM" "218 Xi" "284 ZIA11" "238 ZIA13" "287 ZIc" "235 ZId" "257 ZIr" "236 ZIv")

cd PATH/TO/Atlas/
mkdir WHS_SD_rat_atlas_v4_masks
## create high resolution masks for abovementioned substructures in loop
for pair in "${PAIRS[@]}"
do
set -- $pair
echo "$1 is in $2"
c3d WHS_SD_rat_atlas_v4.nii.gz -thresh "$1" "$1" 1 0 -o
WHS_SD_rat_atlas_v4_masks/"$2"_hrmask.nii.gz
done
```

***Continued***

```
## create unilateral high-resolution mask
cd PATH/TO/Atlas/WHS_SD_rat_atlas_unilateral_masks
for file in PATH/TO/Atlas/WHS_SD_rat_atlas_v4_masks/*_hrmask.nii.gz; do
mask=${file%"_hrmask.nii.gz"}
mask=${mask#"PATH/TO/Atlas/WHS_SD_rat_atlas_v4_masks/"}
echo $mask
if test -f "$mask"_right_hrmask.nii.gz; then
   echo "$mask"_right_hrmask.nii.gz exist
else
         echo "$mask"_right_hrmask.nii.gz does not exist

fslroi $file "$mask"_left_hrmask.nii.gz 0 244 0 1025 0 513
flirt -in "$mask"_left_hrmask.nii.gz -ref 'PATH/TO/Atlas/WHS_SD_rat_atlas_v4.nii.gz' -out
"$mask"_left_hrmask.nii.gz  -applyxfm -dof 6 -nosearch -usesqform
fslmaths $file -sub "$mask"_left_hrmask.nii.gz "$mask"_right_hrmask.nii.gz
fi
done

## create bilateral low-resolution mask
cd ../
mkdir WHS_SD_rat_lr_masks
cd WHS_SD_rat_lr_masks
mkdir bilateral
mkdir unilateral
for file in PATH/TO/Atlas/WHS_SD_rat_atlas_v4_masks/*_hrmask.nii.gz; do
mask=${file%"_hrmask.nii.gz"}
mask=${mask#"PATH/TO/Atlas/WHS_SD_rat_atlas_v4_masks/"}
echo $mask
cp "$file" bilateral/"$mask"_lrmask.nii.gz
fslchpixdim bilateral/"$mask"_lrmask.nii.gz 0.39 0.39 0.39
fslorient -setsform 0.39 0 0 -9.31 0 0.39 0 -24.36 0 0 0.39 -8.88 0 0 0 1 bilateral/"$mask"_lrmask.nii.gz
fslorient -copysform2qform bilateral/"$mask"_lrmask.nii.gz
flirt -in bilateral/"$mask"_lrmask.nii.gz -ref 'PATH/TO/Atlas/atlasX10_downsampled8.nii.gz' -out
bilateral/"$mask"_lrmask.nii.gz -applyxfm -nosearch -usesqform -noresampblur
fslmaths bilateral/"$mask"_lrmask.nii.gz -bin bilateral/"$mask"_lrmask.nii.gz
done
```

***Continued***

```
## create unilateral low-resolution mask
for file in PATH/TO/Atlas/WHS_SD_rat_atlas_unilateral_masks/*_left_hrmask.nii.gz; do
mask=${file%"_left_hrmask.nii.gz"}
mask=${mask#"PATH/TO/Atlas/WHS_SD_rat_atlas_unilateral_masks/"}
echo $mask
cp "$file" unilateral/"$mask"_left_lrmask.nii.gz
fslchpixdim unilateral/"$mask"_left_lrmask.nii.gz 0.39 0.39 0.39
fslorient -setsform 0.39 0 0 -9.31 0 0.39 0 -24.36 0 0 0.39 -8.88 0 0 0 1
unilateral/"$mask"_left_lrmask.nii.gz
fslorient -copysform2qform unilateral/"$mask"_left_lrmask.nii.gz
flirt -in unilateral/"$mask"_left_lrmask.nii.gz -ref 'PATH/TO/Atlas/atlasX10_downsampled8.nii.gz' -out
unilateral/"$mask"_left_lrmask.nii.gz -applyxfm -nosearch -usesqform -noresampblur
fslmaths unilateral/"$mask"_left_lrmask.nii.gz -bin unilateral/"$mask"_left_lrmask.nii.gz
fslmaths bilateral/"$mask"_lrmask.nii.gz -sub unilateral/"$mask"_left_lrmask.nii.gz
unilateral/"$mask"_right_lrmask.nii.gz
done
```
